# The cholesterol biosynthesis enzyme FAXDC2 couples Wnt/β-catenin to RTK/MAPK signaling

**DOI:** 10.1101/2023.12.15.571793

**Authors:** Babita Madan, Shawn Wadia, Siddhi Patnaik, Nathan Harmston, Emile Tan, Iain Bee Huat Tan, W. David Nes, Enrico Petretto, David M. Virshup

## Abstract

Wnts, cholesterol, and MAPK signaling are essential for development and adult homeostasis. Here we report for the first time that fatty acid hydroxylase domain containing 2 (FAXDC2), a previously uncharacterized enzyme, functions as a methyl sterol oxidase catalyzing C4 demethylation in the Kandutsch-Russell branch of the cholesterol biosynthesis pathway. FAXDC2, a paralog of MSMO1, regulates the abundance of specific C4-methyl sterols lophenol and dihydro-TMAS. Highlighting its clinical relevance, FAXDC2 is repressed in Wnt/β-catenin high cancer xenografts, in a mouse genetic model of Wnt activation, and in human colorectal cancers. Moreover, in primary human colorectal cancers, the sterol lophenol, regulated by FAXDC2, accumulates in the cancerous tissues and not in adjacent normal tissues. FAXDC2 links Wnts to RTK/MAPK signaling. Wnt inhibition drives increased recycling of RTKs and activation of the MAPK pathway, and this requires FAXDC2. Blocking Wnt signaling in Wnt-high cancers causes both differentiation and senescence; and this is prevented by knockout of FAXDC2. Our data shows the integration of three ancient pathways, Wnts, cholesterol synthesis, and RTK/MAPK signaling, in cellular proliferation and differentiation.

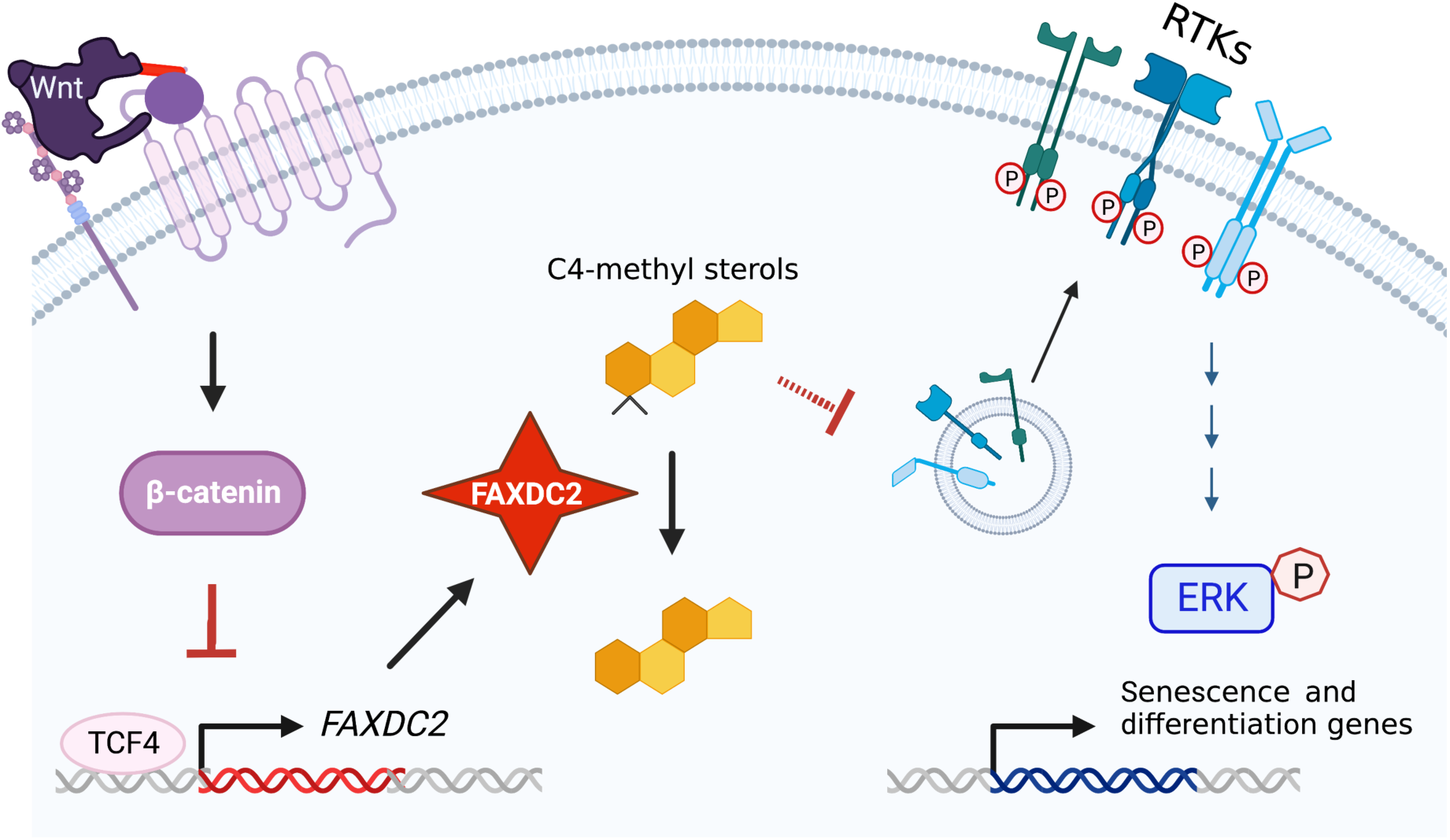

## INTRODUCTION

During animal development and in adult stem cell niches, the balance between stemness, proliferation, and differentiation is regulated by Wnt/β-catenin signaling (1, 2). Wnts are secreted palmitoleated glycoproteins that bind to their cognate receptors on nearby cells to influence multiple downstream signaling pathways in a context-dependent manner. When Wnt signaling is defective, developmental anomalies occur. When Wnt signaling is hyperactive, stem cell differentiation is impaired, and this predisposes to malignancy (3). Studying the pathways regulated by Wnts can reveal both the cellular signaling networks that mediate essential cell fate decisions and the mechanisms of cancer development.

Mutations that cause an accumulation of Wnt receptors on the cell surface (e.g., loss of RNF43 or overexpression of RSPO3) (4–6) lead to hypersensitivity and addiction to Wnts. These mutations enhance Wnt signaling and drive the development of diverse cancers (7). Porcupine (PORCN) is an O-acyltransferase that adds a palmitoleate moiety to all 19 Wnts at a conserved serine residue, an essential step in Wnt ligand secretion (8–10). Pharmacologically targeting PORCN blocks Wnt secretion and prevents the growth of Wnt ligand-dependent cancers. The treatment of Wnt ligand-dependent preclinical xenograft models with drugs that inhibit Wnt palmitoleation provides a powerful system for studying the pathways regulated by Wnt signaling (4, 11–13).

To identify Wnt-regulated genes and pathways, we examined the transcriptional response of multiple Wnt-ligand addicted cancer xenografts following PORCN inhibition (14). Our studies confirmed broad control of transcription by Wnt signaling. In these xenograft models, roughly a third of expressed genes were Wnt-activated, and another third were Wnt-repressed; that is, they had increased expression upon Wnt inhibition. Orthotopic xenograft tumor models identified significantly more Wnt-regulated genes than are found in tissue culture systems (14, 15). This is likely due to the important contribution of the local microenvironment that provides ligands and interactions that are not found in cell culture. For example, these *in vivo* studies have identified ribosomal biogenesis and homologous recombination pathways as Wnt-activated. Recently we also demonstrated that, counterintuitively, Wnt signaling inhibits mitogen-activated protein kinase (MAPK) signaling, a pathway critical for both proliferation and differentiation (16, 17).

Cholesterol biosynthesis is another ancient pathway required for growth. Rapidly proliferating tissues require large amounts of cholesterol for new cell membranes. In addition, cholesterol biosynthesis provides precursors to produce diverse signaling sterols including vitamin D, bile acids, corticosteroids, and sex hormones. Cellular cholesterol is obtained through both diet and *de novo* synthesis. The synthesis of cholesterol from lanosterol proceeds via two parallel pathways, the Bloch and the Kandutsch-Russell (KR) pathways (18–20). Changes in the flux through these pathways alter the abundance of specific cholesterol biosynthesis intermediates, several of which have known biological signaling activity. For example, the C4-dimethyl sterols testicular meiosis activating sterol (T-MAS) and follicular fluid meiosis activating sterol (FF-MAS) regulate germ cell development and EGFR signaling in cancer cells and the C4-monomethyl sterol lophenol regulates cell fate decisions in *C. elegans* (21–23). Individuals with mutations in the genes catalyzing the oxidative removal of C4-methyl groups accumulate these sterols in affected tissues, leading to anomalies such as microcephaly, bilateral congenital cataracts, growth delay, psoriasiform dermatitis, immune dysfunction, and intellectual disability (24–27). Thus, modulation of cholesterol signaling intermediates is critical in development, but whether this is regulated by Wnt signaling is not known.

Here, exploiting potent inhibitors of Wnt production and powerful *in vivo* models, we describe the interaction of Wnt signaling with cholesterol biosynthesis, leading to the regulation of RTK/MAPK signaling. We identify the cholesterol biosynthesis enzyme fatty acid hydroxylase domain containing 2 (FAXDC2) as a key control point. This study establishes the interaction of three ancient and core developmental pathways via cholesterol precursor metabolites, coupling Wnt and RTK/MAPK signaling with broad implications for both normal and cancer biology.

## RESULTS

### Wnt signaling represses the cholesterol biosynthesis enzyme *FAXDC2*

To identify Wnt-regulated genes in cancers, we previously performed a comprehensive transcriptional analysis of multiple Wnt-ligand-dependent models driven by distinct genetic mutations (11, 14, 16). These studies revealed that out of 11,673 expressed genes, inhibiting Wnt signaling decreased the expression of 3,549 (30%) and increased the expression of 4,350 (37%) genes (Fig S1A). One unexpected finding was that Wnt signaling repressed the expression of multiple genes involved in cholesterol biosynthesis (Fig 1A-B).

**Figure 1:**
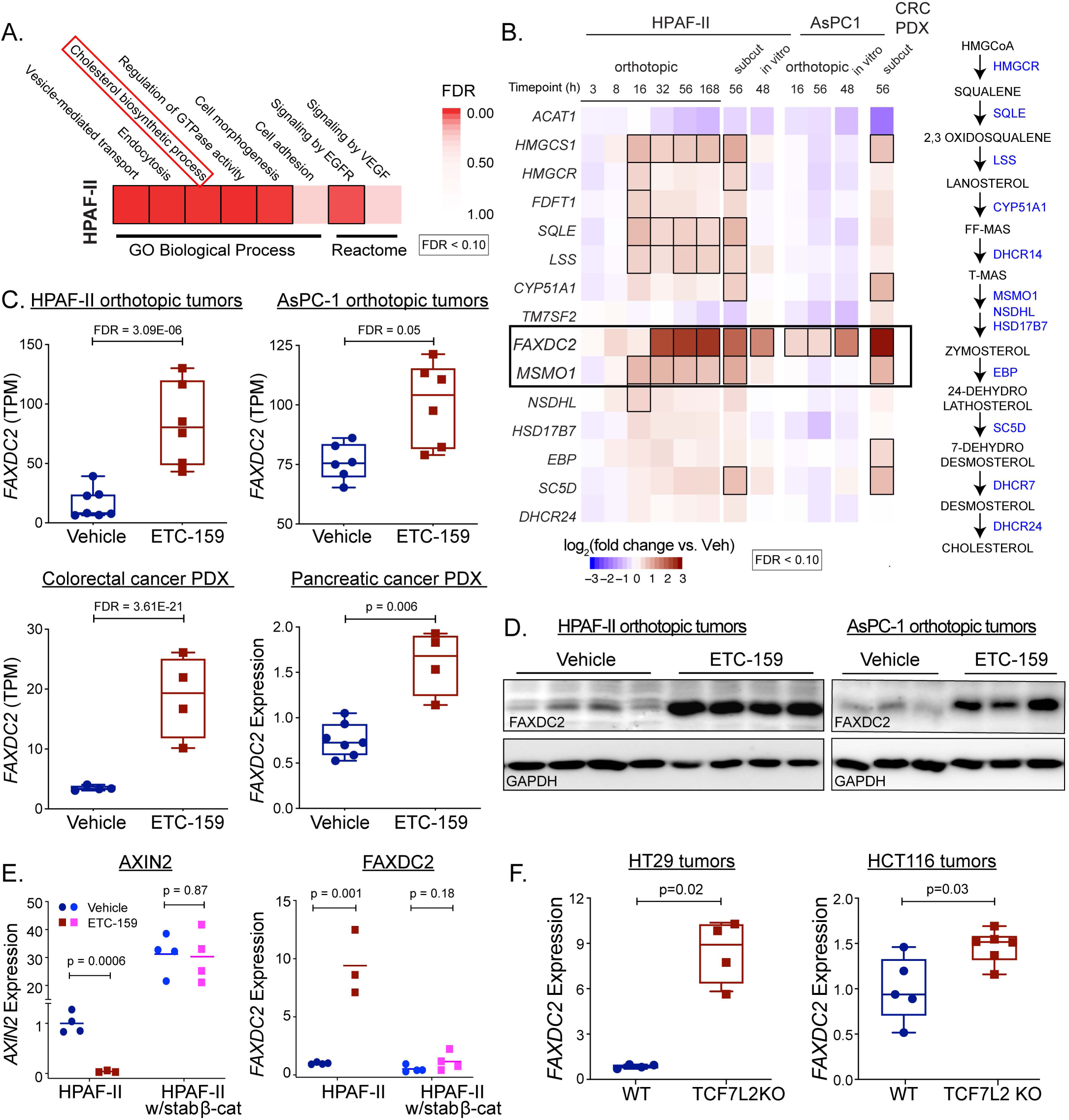
Wnt signaling represses the cholesterol biosynthesis enzyme FAXDC2. **A. *Wnt inhibition upregulates genes regulating cholesterol biosynthesis.*** Pharmacologic Wnt inhibition with ETC-159 in HPAF-II orthotopic xenografts upregulates 4,350 genes that are grouped into temporal clusters (n=4-6 mice/group). GO Biological Process and Reactome analysis of the *Wnt-repressed* genes highlight processes including cholesterol biosynthesis, vesicle-mediated transport, and EGFR/ VEGF signaling (hypergeometric test, FDR < 10%). B. ***Wnt signaling represses genes encoding enzymes in the cholesterol biosynthesis pathway***: Left, Wnt inhibition with ETC-159 increases expression of multiple cholesterol biosynthesis pathway genes in three independent systems: HPAF-II and AsPC-1 orthotopic xenografts and cells in culture as well as a colorectal PDX (14) (n = 4-6 tumors/group). Boxed genes show significant change in expression (FDR < 0.10). Right, cholesterol biosynthesis pathway. C. ***Wnt inhibition increases FAXDC2 expression in multiple Wnt-addicted cancer models:*** ETC-159 treated HPAF-II, AsPC-1 orthotopic tumors, and Wnt-addicted colorectal and pancreatic cancer patient-derived xenograft models have 2-10 fold higher *FAXDC2* expression compared to the control tumors. Relative expression from RNA seq in TPM or RT-qPCR is shown. Each data point represents an independent tumor, n = 5-6/group. D. ***ETC-159 treatment increases FAXDC2 protein abundance in tumors*.** Protein lysates from the HPAF-II and AsPC-1 orthotopic xenografts from the vehicle or ETC-159 treated mice were probed with FAXDC2 or GAPDH antibodies. Each lane represents tumor lysate from an individual mouse. E. ***Stabilized β-catenin suppresses FAXDC2 expression despite upstream Wnt inhibition by ETC-159***. Mice bearing xenografts from control HPAF-II cells or cells expressing stabilized β-catenin were treated with ETC-159 or vehicle for 56 h. *FAXDC2* and *AXIN2* mRNA were quantitated by qRT-PCR and normalized to both *ACTB* and *EPN1. AXIN2* upregulation is a control for stabilized β-catenin activity. Each data point represents an independent tumor. F. Genetic inhibition of Wnt signaling by TCF7L2 knockout in HT-29 and HCT116 colon cancer xenografts increases FAXDC2 expression by two to eight-fold compared to the WT controls. Each data point represents an independent tumor.

The gene in the cholesterol biosynthesis pathway with the most significant and general upregulation after inhibition of Wnt signaling was Fatty Acid Hydroxylase Domain Containing 2, *FAXDC*2 (C5orf4) (Figure 1B), a putative paralog of methyl sterol monooxygenase 1 (MSMO1) about which little is known. Treatment of mice bearing HPAF-II orthotopic tumors with a pan-Wnt secretion inhibitor ETC-159 led to a robust upregulation of *FAXDC2* mRNA (7.5-fold after 2 days of treatment) (Fig 1B-C). *FAXDC2* expression was also markedly enhanced by Wnt inhibition in four independent Wnt-high cancer models driven by distinct genetic mutations. These included AsPC-1 pancreatic cancer orthotopic xenografts and pancreatic cancer patient-derived xenograft (PDX) PAXF1861, both with loss-of-function RNF43 mutations, and a colorectal PDX with an R-spondin translocation, (Fig 1B-C). This confirms that regulation of *FAXDC2* by Wnts is widespread. Consistent with the change in gene expression, we found a marked increase in FAXDC2 protein abundance in both Wnt-inhibited HPAF-II and AsPC-1 orthotopic tumors (Fig 1D).

Wnt pathway inhibition causes phosphorylation-mediated degradation of β-catenin. To test if degradation of β-catenin is required for the robust upregulation of *FAXDC2* and other genes in the cholesterol biosynthesis pathway, we generated HPAF-II cells that express stabilized β-catenin (16). β-catenin with mutated serine residues (S4A: S33A, S37A, T41A, and S45A) is insensitive to CK1α/GSK3 phosphorylation, preventing its proteasomal degradation. HPAF-II tumors expressing stabilized β-catenin have sustained downstream activation of the Wnt/β-catenin pathway regardless of PORCN inhibition, as evidenced by the failure of PORCN inhibition to reduce the expression of the Wnt target gene *AXIN2* (Fig 1E, left). Stabilized β-catenin prevented the upregulation of several cholesterol pathway genes. Notably, the gene with the highest dynamic range was FAXDC2 (Fig 1E, S1B). Thus, active β-catenin represses *FAXDC2* expression.

To complement the PORCN inhibitor assays, we performed a genetic test of the role of β-catenin signaling in suppressing *FAXDC2* expression. We examined the impact of knocking out β-catenin’s nuclear binding partner, TCF4, which is encoded by the *TCF7L2* gene (28). APC-mutant HT29 and β-catenin mutant HCT116 colorectal cancer cells with or without *TCF7L2* knockout were obtained from the Hecht group and xenograft tumors were generated (29). *TCF7L2* knockout tumors had, as previously demonstrated, decreased β-catenin signaling as evidenced by reduced *AXIN2* expression (Fig S1C-D). Notably, both HT29 *TCF7L2* KO and HCT116 *TCF7L2* KO xenograft tumors showed ∼2-10-fold increase in the expression of *FAXDC2* (Fig 1F). These data establish that the Wnt-β-catenin-TCF4 axis strongly represses *FAXDC2* expression.

SREBP1 and SREBP2 are well-established regulators of several genes in the cholesterol biosynthesis pathway. To directly test the impact of SREBP1 and SREBP2 on *FAXDC2* expression, we used multiple siRNAs to knock down these genes in HPAF-II cells. We find that knockdown of neither SREBP1 nor SREBP2 affected Wnt inhibition induced increase in *FAXDC2* expression (Fig S1E-F). We conclude that SREBP regulates neither basal nor Wnt-regulated FAXDC2 expression.

### FAXDC2 is a C4-methyl sterol oxidase in the Kandutsch-Russell branch of the cholesterol biosynthesis pathway

During cholesterol biosynthesis, squalene is oxidized and cyclized to form lanosterol, a C4-dimethyl sterol (Fig 2A-B). Conversion of lanosterol to cholesterol proceeds down two parallel routes known as the Bloch and Kandutsch-Russell (KR) pathways and involves nine distinct enzymes (18, 30, 31) (Fig 2A). One of the key steps in this process is the removal of the two methyl groups at the C4 position. The oxidative removal of these C4 methyl groups in the endoplasmic reticulum is catalyzed in a stepwise series of reactions (Fig 2A-B) by methyl sterol monooxygenase (MSMO1), NAD(P) dependent steroid dehydrogenase-like (NSDHL) and keto-reductase hydroxysteroid 17-beta dehydrogenase 7 (HSD17B7) (32, 33).

**Figure 2:**
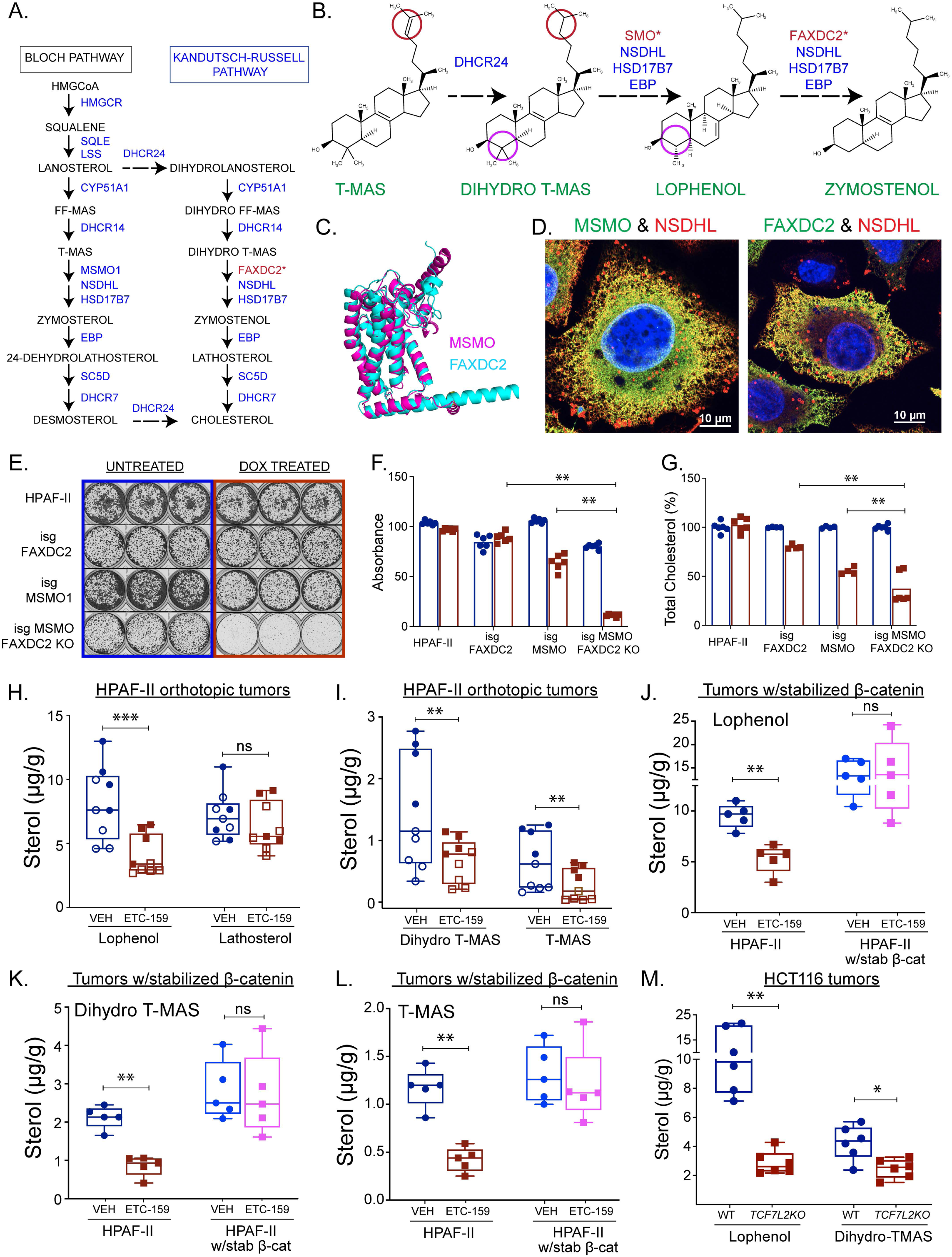
FAXDC2 is a C4-demethylase in the Kandutsch-Russell branch of the cholesterol biosynthesis pathway. A. ***Diagram illustrating post-lanosterol cholesterol biosynthesis*** via Bloch and Kandutsch-Russell pathways, highlighting key steps and associated enzymes. *Proposed location of FAXDC2 in the pathway. B. Diagram depicting the conversion of T-MAS to Zymostenol, highlighting reduction at C24 and demethylation at C4 involving sterol methyl oxidase (SMO). *SMO indicates either MSMO1 or FAXDC2 sterol methyl oxidase. C. MSMO and FAXDC2 structures predicted by AlphaFold show considerable homology (root-mean-square deviation of atomic positions 1.36 Å). D. ***Like MSMO1, FAXDC2 colocalizes with NSDHL in the SER***: Epitope-tagged MSMO1, NSDHL, and FAXDC2 constructs were expressed in HeLa cells and their localization was visualized by staining with fluorescence-tagged anti-epitope antibodies. **E-G. *Combined knockdown of FAXDC2 and MSMO1 reduces total cholesterol levels and prevents cellular proliferation***: HPAF-II cells, HPAF-II cells with doxycycline (dox)-inducible single guide RNAs (isg) targeting *FAXDC2* or *MSMO1* alone, and *FAXDC2* KO cells with dox-isg targeting *MSMO1* were cultured for ten days. (E) Representative images of crystal violet staining from three independent experiments. (F) Crystal violet dye was solubilized, and absorbance was measured at 570 nm. (G) Total cholesterol levels were significantly reduced in the FAXDC2 and MSMO1 double KO cells compared to the single knockouts. **H-I. *ETC-159 treatment reduces C4-methyl sterols in HPAF-II orthotopic tumors with high FAXDC2 expression***. Sterol levels in mice treated with vehicle or ETC-159 were measured by GC-MS. Data from individual tumor samples (n= 4-5/group) from two independent experiments were combined to calculate p values (open and closed symbols), controlling for batch effects. **J-L. *Stabilized β-catenin, which represses FAXDC2, increased C4-methyl sterol levels independent of upstream Wnt signaling inhibition***. C4-methyl sterol levels in HPAF-II tumors with and without stabilized β-catenin from mice treated with ETC-159 or vehicle were analyzed by GC-MS. Each point denotes an individual tumor. n = 4-5/group. **M**. ***C4-methyl sterols are reduced in TCF7L2 KO HCT116 xenografts compared to the control tumors.*** Sterol levels were measured by GC-MS. Each point depicts an individual tumor sample.

FAXDC2 is annotated as a paralog of MSMO1 (Alliance of Genome Resources HGNC:1334, Version: 5.3.0) (34) but has not been extensively studied. Consistent with the annotation, FAXDC2 has 27.5% amino acid identity and 49% similarity with MSMO1 and 24.2% identity with another C4 methyl sterol oxidase, ERG25 in *Saccharomyces cerevisiae* (Fig S2A). All contain the three conserved histidine-rich motifs (HX3H, HX2HH, and HX2HH) characteristic of membrane-bound non-heme iron oxygenases (35). Orthologs of FAXDC2 are similarly conserved in plants and sponges (Fig S2A) (36, 37). The Alphafold-predicted structures (38) of MSMO1 and FAXDC2 are nearly superimposable with an RMSD of 1.4 Å (Fig 2C). Finally, previous studies in plants show that two orthologs of MSMO1 cooperate with NSDHL and other enzymes critical for the demethylation of cholesterol biosynthesis intermediates (CBIs). Consistent with FAXDC2 being a paralog of MSMO1 and acting as a C4-demethylase, immunofluorescence experiments to interrogate the interactions of enzymes of the demethylase complex show that FAXDC2, like MSMO1, can co-localize with NSDHL in the smooth ER (Fig 2D and S2B). Thus, sequence homology, predicted structure, and intracellular localization are all consistent with FAXDC2 being a C4-methyl sterol oxidase.

If FAXDC2 is a functional paralog of MSMO1 in the cholesterol biosynthesis pathway, cells lacking either MSMO1 or FAXDC2 should have reduced cholesterol synthesis and impaired cell growth in media with delipidated cholesterol-poor serum. Moreover, the knockout of both enzymes should turn off both KR and Bloch arms of the cholesterol biosynthesis pathway and further compromise cell growth. *FAXDC2* knockout alone in HPAF-II cells (where expression of *FAXDC2* is already repressed/low) did not reduce cell viability as assessed by colony formation (Fig 2E-F). In contrast, inducible knockout of *MSMO1* caused a ∼40% decrease in colony formation, suggesting that these cells were more reliant on flux through the Bloch pathway. Notably, inducible knockout of *MSMO1* in cells lacking *FAXDC2* led to a ∼90% decrease in colony-forming ability. Under these conditions, total cellular cholesterol levels were markedly decreased by *MSMO1* knockout and further reduced in cells lacking both *MSMO1* and *FAXDC2* (Fig 2G). This is consistent with FAXDC2 being a paralog of MSMO1 in the cholesterol biosynthesis pathway.

### FAXDC2 regulates the abundance of C4-methyl sterols

To test the function of FAXDC2 as a C4-methyl sterol oxidase involved in cholesterol biosynthesis, we postulated that there should be differences in sterol composition between the control FAXDC2-low (Wnt-high) and ETC-159 treated, FAXDC2-high (Wnt-low) HPAF-II tumors. To test this, we measured the abundance of cholesterol biosynthesis intermediates (CBIs) in the vehicle and ETC-159 treated tumors from two independent experiments. GC-MS analysis identified three C4-methyl sterols in FAXDC2-low HPAF-II orthotopic xenografts: 4α-methylcholest-7-enol (lophenol), 4,4-dimethylcholest-8-enol (dihydro-TMAS), and 4,4-dimethylcholesta-8,24-dienol (TMAS) (Fig S3A-B) (18). The abundance of all three C4 methyl sterols significantly decreased as FAXDC2 abundance increased in the tumors treated with the Wnt inhibitor (Fig 2H-I). These biochemical data suggest that FAXDC2 catalyzes a rate-limiting C4-demethylation of specific cholesterol biosynthesis intermediates. High levels of FAXDC2 can drive enhanced flux through the cholesterol biosynthesis pathway and hence reduce the abundance of lophenol, dihydro-TMAS, and TMAS, consistent with FAXDC2 being a methyl sterol oxidase.

Our model predicts that ectopically stabilized β-catenin, by repressing FAXDC2 expression (Fig 1E), should block the ETC-159-induced changes in cholesterol biosynthesis and prevent changes in the abundance of C4-methyl sterols. Consistent with this, tumors with stabilized β-catenin had low FAXDC2 (Fig 1E) and higher basal levels of lophenol and dihydro-TMAS than the HPAF-II tumors (Fig 2J-L). Furthermore, the abundance of these C4-methyl sterols was unaltered by ETC-159 treatment in HPAF-II tumors with stabilized β-catenin, confirming that the drug acts on cholesterol biosynthesis through its effects on the Wnt/β-catenin pathway.

In HCT116 colorectal cancer xenografts that have hyperactive β-catenin and low *FAXDC2*, we also observed accumulation of C4-methyl sterols lophenol and dihydro TMAS. In these xenografts, genetic inhibition of β-catenin activity by TCF7L2 KO also significantly reduced the abundance of the sterols, again inversely correlated with *FAXDC2* expression (Figs 1F and 2M).

The two C4-methyl sterols that are most robustly decreased upon Wnt inhibition, lophenol and dihydro-TMAS, being saturated at C24 (Fig. 2B), are specific to the Kandutsch-Russell (KR) arm of the cholesterol biosynthesis pathway. The key difference between the Bloch and the KR pathway is the step at which C24 double bond is reduced by 24-dehydrocholesterol reductase (DHCR24). The Bloch pathway utilizes DHCR24 in the last step of cholesterol biosynthesis, while the KR pathway uses DHCR24 to convert lanosterol to 24,25-dihydro-lanosterol much earlier in the biosynthesis of cholesterol (Fig 2A). This flow is largely unidirectional such that the Bloch pathway intermediates may crossover into the KR pathway at multiple steps but not necessarily *vice versa* (31). The correlation of changes in 24, 25-dihydro C4-methyl sterols in HPAF-II xenografts with FAXDC2 levels suggests that FAXDC2 is a methyl sterol oxidase predominantly in the KR pathway and that it catalyzes the steps between dihydro-T-MAS and zymostenol, parallel to the activity of MSMO1 in the Bloch pathway (Fig 2A-B). Low FAXDC2 activity in Wnt-high tumors causes an accumulation of upstream C4-methyl sterols of the KR pathway. In contrast, an increase in FAXDC2 activity in Wnt-inhibited tumors enhances flux through this branch of the pathway, leading to the depletion of C4-methyl sterols (lophenol and dihydro-TMAS) upstream of the steps it catalyzes (Fig 2H-I).

Taken together, these data show that FAXDC2 is a C4-methyl sterol oxidase that is repressed by high Wnt/β-catenin signaling. FAXDC2 acts preferentially in the KR branch of the cholesterol biosynthesis pathway, and its upregulation changes flux through the pathway, thereby altering the abundance of specific C4-methyl sterols.

### *FAXDC2* is repressed in Wnt-high human cancers and leads to an accumulation of C4-methyl sterols

Pathologic Wnt activation occurs in over 80% of human colorectal cancers. To examine if *FAXDC2* was repressed in Wnt-high cancers, we analyzed the data from Gene Expression Profiling Interactive Analysis (GEPIA 2.0) (39). Consistent with our xenograft data, *FAXDC2* expression was significantly repressed in human colorectal cancers compared to normal colorectal tissue (Fig 3A). Additionally, promoter analysis using datasets from ENCODE (40) showed that the H3K4me3 activation mark was present on the promoter of *FAXDC2* in normal colon tissue but was completely absent in two colon cancer cell lines (Fig 3B).

**Figure 3:**
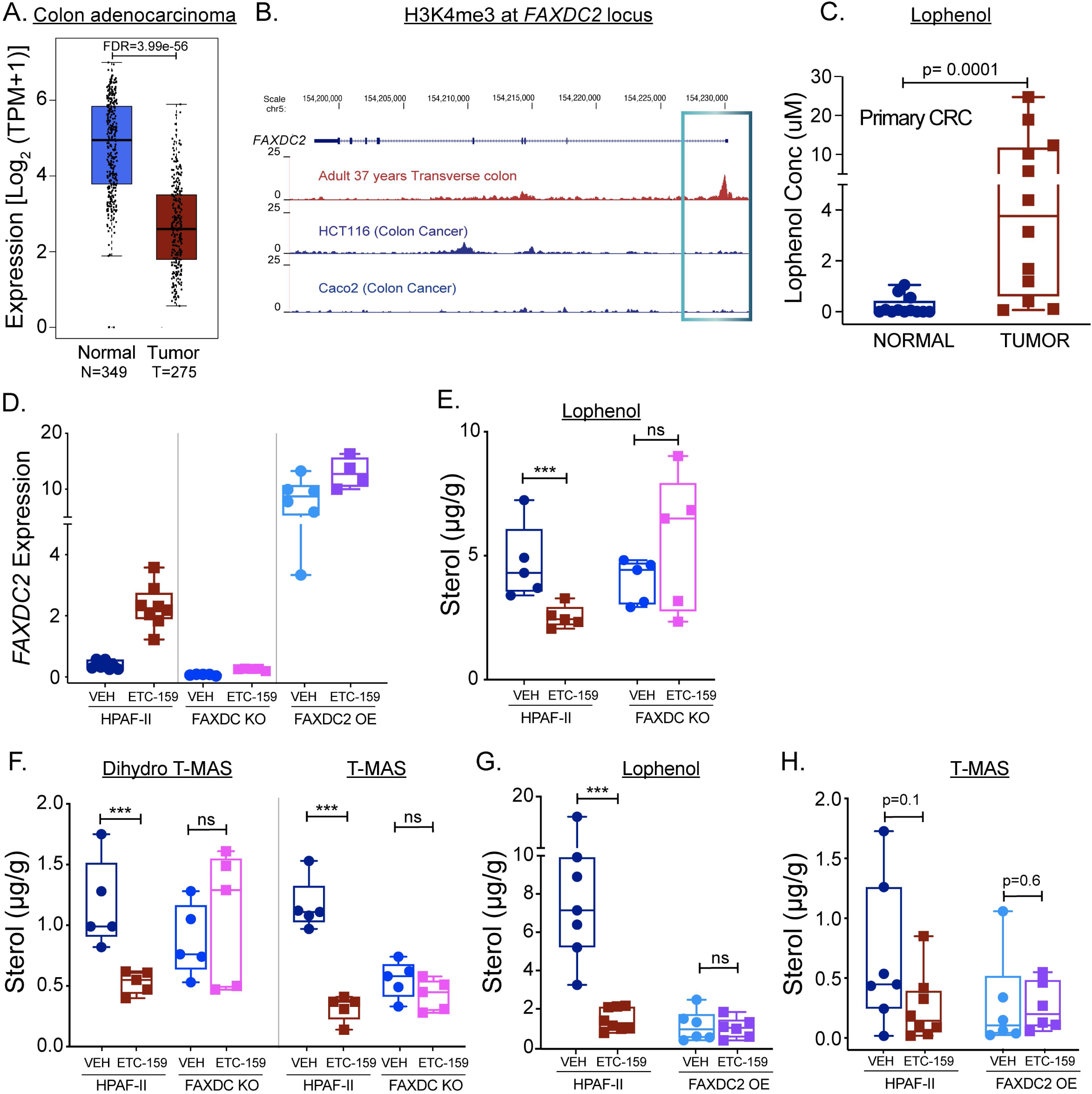
*FAXDC2* expression correlates with and drives changes in C4-methyl sterols. ***A-C.* Primary Wnt-high colorectal cancers have repressed FAXDC2 expression and high levels of lophenol** A. ***FAXDC2 is repressed in primary colorectal cancers compared to corresponding normal tissues:*** Expression of *FAXDC2* was compared in the normal tissues vs. tumor tissues using GEPIA 2.0 (TCGA and GTEx datasets). B. **H3K4me3**, a marker of active transcription, marks the *FAXDC2* genomic locus in the normal colon but is absent in cancer cell lines. Data from Cistrome DB. C. ***Lophenol is markedly elevated in primary colorectal cancers***. Sterol abundance was measured in primary colorectal cancers and adjacent normal tissue by GC-MS. Each point represents lophenol levels in an individual tumor, n=12 tumors/group. **D-H. Manipulation of FAXDC2 expression regulates C4-methyl sterol abundance.** D. Relative expression of *FAXDC2* in HPAF-II, FAXDC2 KO, and FAXDC2 OE xenografts as measured by qRT-PCR. **E-F. *Knockout of FAXDC2 abrogates the ETC-159 treatment-induced change in the abundance of C4-methyl sterols***. Sterol abundance was measured in HPAF-II and FAXDC2 KO xenografts from control and ETC-159 treated mice. Each point represents the level of indicated methyl sterols measured using GC-MS in an individual tumor sample. n= 4-5/group. **G-H. *FAXDC2 overexpression reduces lophenol and T-MAS levels comparable to the Wnt-inhibited tumors***. Sterol abundance was measured in HPAF-II and FAXDC2 overexpressing xenografts from control and ETC-159 treated mice. Each point represents the abundance of methyl sterols measured using GC-MS in an individual tumor sample. n= 4-5/group.

If *FAXDC2* is repressed in most human colorectal cancers, then methyl sterol intermediates should accumulate in those cancers. To test this, we measured both lophenol abundance and *FAXDC2* expression in locally available matched normal and tumor tissues from Wnt-high colorectal cancer patients. Consistent with the GEPIA data, in fifty paired normal and tumor samples from colorectal cancer patients, *FAXDC2* expression was significantly lower in the tumors compared to the normal tissues (Fig S4A). Most strikingly, these colorectal cancers had markedly elevated levels of lophenol compared to its low abundance in the matched normal tissues with high *FAXDC2* levels (Fig 3C). These data are similar to our findings in Wnt-high pancreatic cancer xenografts and support the model that elevated Wnt/β-catenin signaling represses *FAXDC2* expression, decreasing flux through the Kandutsch-Russell branch of the cholesterol biosynthesis pathway leading to the accumulation of upstream intermediates, including lophenol and other C4 methyl sterols.

### FAXDC2 mediates Wnt-regulated changes in C4-methyl sterols

Our data show a strong inverse correlation between FAXDC2 levels and C4-methyl sterol abundance. To directly test if FAXDC2 is required for the changes in sterol abundance, we used Cas9/CRISPR to create FAXDC2 knockout clones using sgRNAs (Fig S4B-C). HPAF-II WT or *FAXDC2* KO cells were injected into mice to generate xenograft tumors. The absence of FAXDC2 transcript and protein in the KO tumors was confirmed by qRT-PCR and immunoblots (Fig 3D and S4D). *FAXDC2* expression and C4-methyl sterol abundance were measured before and after seven days of Wnt inhibition. Knockout of *FAXDC2* abrogated the increase in *FAXDC2* expression induced by Wnt inhibition (Fig 3D). Notably, compared to the control tumors, *FAXDC2* KO blocked the Wnt inhibition-mediated decrease in the C4-methyl sterols of the KR pathway, lophenol, and dihydro-TMAS (Fig 3E-F). This shows directly that the absence of FAXDC2 causes an accumulation of C4-methyl sterols; hence it is required for their demethylation.

Next, we rescued the expression of *FAXDC2* in the KO cells by stably expressing it under the control of a CMV promoter (hereafter referred to as *FAXDC2* OE cells). The xenografts from these cells had a ten-fold higher than normal expression of *FAXDC2* (Fig 3D). As predicted, the *FAXDC2* OE xenografts from the vehicle-treated mice had markedly reduced levels of lophenol and T-MAS, comparable to what was seen following Wnt inhibition in HPAF-II tumors when FAXDC2 levels increased (Fig 3G-H). Notably, *FAXDC2* OE tumors showed no further decrease in sterol abundance following Wnt inhibition, directly demonstrating that FAXDC2 regulates flux through the cholesterol biosynthesis pathway and mediates Wnt-regulated changes in C4-methyl sterol abundance. Taken together, these studies establish that FAXDC2 is a Wnt-repressed enzyme in the KR branch of the cholesterol biosynthesis pathway that regulates the abundance of 24-saturated C4-methyl sterols.

### FAXDC2 activity influences Wnt-regulated gene expression

While acting as precursors for cholesterol, C4-methyl sterols also act as signaling molecules in major biological processes, including auxin transport in plants, dauer formation in *C.elegans*, oogenesis, and oligodendrocyte differentiation in mammals (22, 41, 42). We speculated that FAXDC2-regulated changes in C4-methyl sterols might also regulate downstream signaling pathways. To test this, we assessed the transcriptional landscape of HPAF-II xenografts with or without *FAXDC2* knockout (Fig 4A, column 1). Knockout of *FAXDC2* altered the expression of 3,570 genes compared to the parental HPAF-II tumors (fold change>1.5, FDR<10%), consistent with a significant role for C4-methyl sterols in cellular signaling pathways. These FAXDC2 dependent genes were grouped into 6 clusters (Fig 4A, S5A-B).

**Figure 4:**
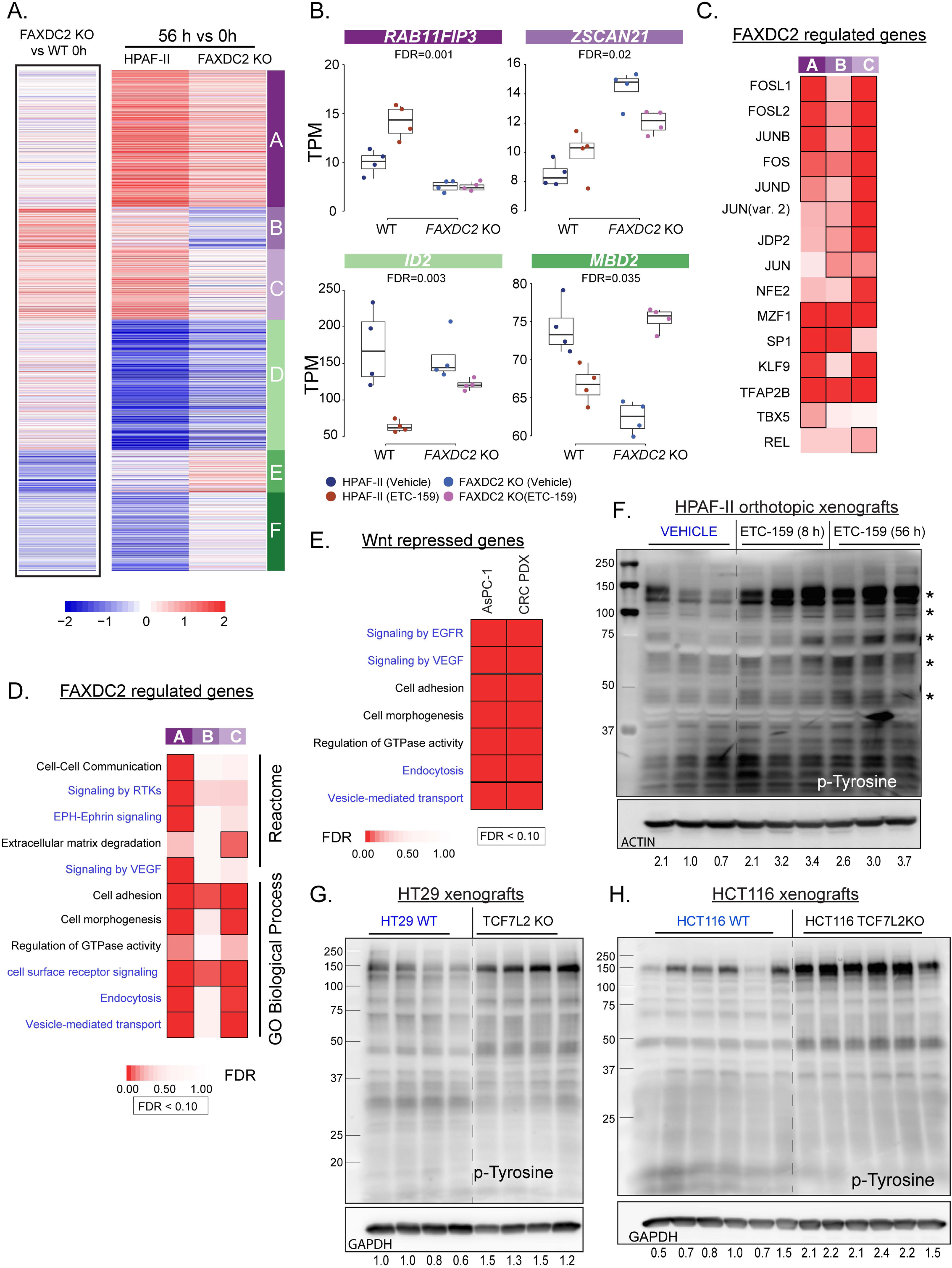
FAXDC2 is a downstream effector of Wnt signaling, and *FAXDC2-regulated* genes are enriched for RTK signaling. **A. *The expression of a subset of Wnt-regulated genes depends on FAXDC2 (FDR < 10%).*** Heatmap of the fold changes of 3560 *FAXDC2*-dependent Wnt-regulated genes (interaction test, FDR<10%). Left panel shows gene expression changes (log2 fold changes) between WT and FAXDC2 KO tumors at 0h. Right panel shows gene expression changes following treatment with ETC-159 in both parental HPAF-II tumors (56 h vs 0 h) and FAXDC2 KO tumors (56 h vs 0 h). This set of differentially responding genes was partitioned into six distinct clusters based on similarities in their response to ETC-159, including three clusters of Wnt-repressed genes (A-C) and three clusters of Wnt activated genes (D-F). **B.** Representative genes from four clusters in figure 4A show that FAXDC2 KO blunts or reverses the response to Wnt inhibition (TPM - transcripts per million). **C.** FAXDC2-regulated Wnt-repressed genes in clusters A-C show an enrichment of AP1 family TFBS motifs in their promoters (FDR < 10%). **D. *RTK signaling pathways are enriched in FAXDC2-regulated Wnt-repressed genes.*** Data from representative GO Biological Process and Reactome enrichment of Wnt-repressed genes from clusters A-C that were identified in Figure 4A (hypergeometric test, FDR < 10%). **E.** GO Biological Processes and Reactome enrichment analysis of Wnt-repressed genes in ETC-159 treated AsPC-1 and CRC PDX tumors shows an upregulation of RTK signaling pathways (hypergeometric test, FDR < 10%). **F. *Pharmacologic Wnt inhibition with ETC-159 in HPAF-II xenografts increases protein tyrosine phosphorylation.*** HPAF-II orthotopic tumor protein lysates from the vehicle or ETC- 159 treated mice were separated on a 10% SDS gel. The blots were probed with phosphotyrosine antibodies. Each lane represents tumor lysate from an individual mouse. **G-H**. ***Genetic inhibition of Wnt/β-catenin signaling in TCF7L2 KO HT29 and HCT116 xenografts increases protein tyrosine phosphorylation.*** Protein lysates from WT and *TCF7L2 KO* HT29 and HCT116 tumors were analyzed for abundance of phosphotyrosine proteins as in F. Each lane contains tumor lysate from an individual mouse.

As noted, Wnt inhibition regulates the expression of 11,673 genes in HPAF-II orthotopic xenografts (Fig S1A). The response of 2,159 of the Wnt-regulated genes in HPAF-II xenografts depended on FAXDC2. FAXDC2 regulated 1,146 Wnt-repressed genes, i.e., genes that were upregulated by Wnt inhibition in the HPAF-II WT tumors but were unaltered in FAXDC2 KO tumors (clusters A-C Fig 4A, Table S1) (FDR<10%, interaction test) and 1,013 Wnt-activated genes (Clusters D-F Fig 4A). Examples of such FAXDC2-regulated genes, *RAB11FIP3*, *IGFBP3, ID2*, and MBD2, are shown in Fig 4B and S5C. Thus, FAXDC2 is essential for regulating a significant proportion of the genes whose expression changes upon Wnt inhibition.

To better understand the signaling pathways regulated by FAXDC2, we performed transcription factor binding site analysis of the FAXDC2-regulated genes and found enrichment for ETS and AP1 binding sites (Fig 4C, Table S2). ETS and AP1 transcription factors act downstream of MAPK signaling to regulate gene transcription, suggesting a role for FAXDC2 in regulating MAPK signaling. This is consistent with our previous reports that Wnt inhibition activates MAPK signaling (16, 17). Gene Ontology (GO) and Reactome pathway analysis of these FAXDC2-dependent genes also showed significant enrichment for genes regulating vesicle-mediated transport, signaling by Receptor Tyrosine Kinases (RTKs), and cell-cell communication processes (Fig 4D). A similar enrichment was also present in Wnt-regulated genes in all three Wnt-ligand addicted models, HPAF-II and AsPC-1 orthotopic xenografts and CRC PDX (Figs 1A and 4E). This is consistent with FAXDC2 being a major downstream effector of Wnt signaling and suggests that Wnt signaling, via FAXDC2, could regulate RTKs and downstream MAPK signaling.

### Wnt inhibition leads to an increase in Receptor Tyrosine Kinase activation in Wnt-high tumors

To examine if Wnt inhibition activated protein tyrosine kinases, we analyzed changes in tyrosine phosphorylation in the vehicle and ETC-159 treated HPAF-II tumor xenografts. Remarkably, Wnt inhibition led to a robust increase in phosphotyrosine staining of multiple proteins in tumor lysates (Fig 4F). The increase in tyrosine phosphorylation of multiple proteins in response to Wnt inhibition was confirmed by a phosphotyrosine array (Fig S5D). This shows that tyrosine kinase signaling is activated upon Wnt inhibition in these tumors. Similarly, genetic inhibition of Wnt/β-catenin signaling (by knockout of *TCF7L2*) caused enhanced tyrosine phosphorylation in two additional models, APC-mutant HT29 and β-catenin mutant HCT116 colorectal cancer xenografts (Fig 4G-H). Thus, inhibiting Wnt/β-catenin signaling either pharmacologically or genetically leads to the activation of cellular tyrosine phosphorylation.

### Wnt inhibition increases the recycling of Receptor Tyrosine Kinases

RTK protein abundance and activity is regulated by a balance of endocytosis, recycling, and degradation (43, 44). Gene ontology and reactome analysis indicates that Wnt signaling regulates receptor recycling. Reduced endocytosis and lysosomal degradation lead to increased cell surface abundance of RTKs, where they are available for interaction with their ligands and activation of downstream signaling. We therefore tested if Wnt inhibition increased the cell surface abundance of EGFR, a prototype for RTK endocytic recycling. Indeed, the abundance of EGFR on the surface of cultured Wnt-ligand-dependent pancreatic cancer cell lines HPAF-II and AsPC-1 increased significantly following Wnt inhibition (Fig 5A and S6A). Likewise, a clear re-localization of EGFR to the cell membrane following ETC-159 treatment was seen by indirect immunofluorescence staining of non-permeabilized HPAF-II cells (Fig 5B). These data indicate that enhanced EGFR recycling to the plasma membrane upon Wnt inhibition is a general phenomenon in Wnt-ligand-dependent cancers.

**Figure 5:**
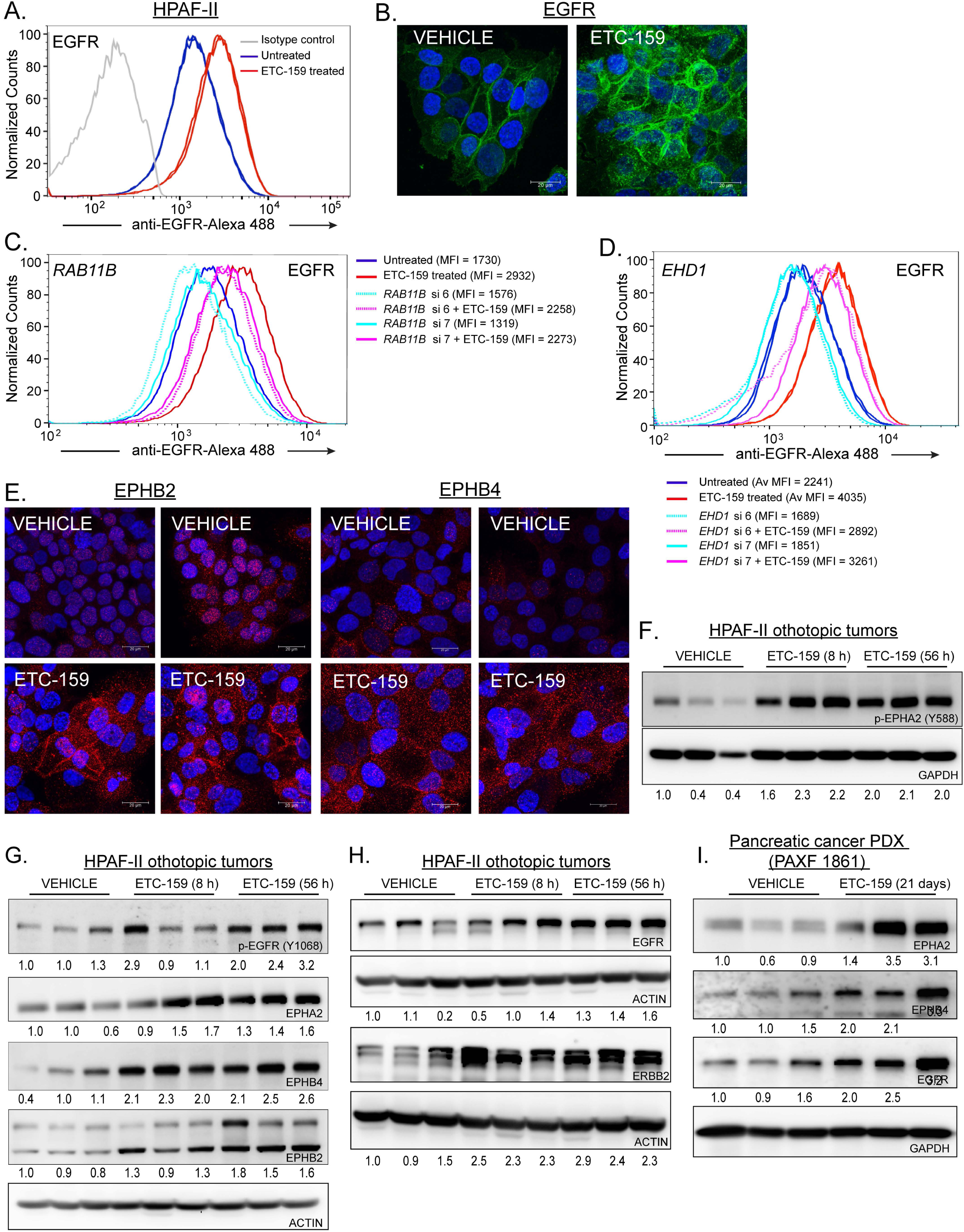
Increased RTK recycling in Wnt-addicted cancers upon Wnt inhibition. **A-B. *ETC-159 treatment of HPAF-II cells increases abundance of EGFR on the cell surface***. HPAF-II cells were treated with 100 nM ETC-159 for 72 h. **(A)** Cells were stained with Alexafluor-488 conjugated anti-EGFR antibody and analyzed by flow cytometry. Av. MFI = Average median fluorescence intensity of the technical replicates from the same experiment. Each histogram represents ∼50,000 cells. Data is representative of three independent experiments (p = 0.009). **(B)** Endogenous EGFR levels on non-permeabilized cells were assessed by indirect immunofluorescence microscopy. **C-D*. Partial knockdown of RAB11B or EHD1 blunts the ETC-159-induced EGFR increase on the surface of HPAF-II cells.*** HPAF-II cells were transfected with two independent siRNAs (si 6 and si 7) against *RAB11B* (C) *or EHD1* (D) for 24 h, followed by treatment with ETC-159 for 72 h. EGFR levels on the cell surface were assessed by flow cytometry. Each histogram represents ∼50,000 cells from one replicate. Data is representative of three independent experiments. Average MFI of the technical replicates is shown. D. ***EPHB2 and EPHB4 levels are increased by Wnt inhibition***. HPAF-II cells treated with 100 nM ETC-159 for 72 h, and the levels of endogenous EPHB2 and EPHB4 on non-permeabilized cells were assessed by indirect immunofluorescence microscopy. **F-I. *Wnt inhibition increases activation of EPHA2 and EGFR receptor tyrosine kinases and the protein abundance of multiple receptor tyrosine kinases in HPAF-II xenografts and pancreatic PDX models***. Protein lysates from HPAF-II orthotopic xenografts or pancreatic PDX from vehicle or ETC-159 treated mice were analyzed for the expression of p-EphA2, p-EGFR and abundance of indicated RTKs by western blots. Each lane represents tumor lysate from an individual mouse. (H) The protein lysates were prepared as a master mix and loaded on independent gels. Only one blot was probed for the load control which is shared with Fig 4F.

To further assess the role of endocytic recycling in regulating Wnt inhibition-mediated changes in cell surface EGFR abundance, we knocked down multiple genes associated with recycling endosomes (45, 46). While knockdown of RAB4, RAB25, and RAB11A had no effect, knockdown of *RAB11B* using two independent siRNAs substantially reduced the ETC-159 mediated increase in EGFR levels on the surface of HPAF-II cells (Fig 5C and S6B-H) (47, 48). Similar to *RAB11B*, the knockdown of *EHD1* using two independent siRNAs also mitigated the ETC-159 mediated increase in the cell surface abundance of EGFR in HPAF-II cells (Fig 5D and S6I) (49). Taken together, these data suggest that enhanced late endocytic recycling leads to an increase in cell surface abundance of EGFR following Wnt inhibition.

Rab7 is a key Rab required for the transport from late endosomes to lysosomes. As can be seen in Fig S6J-K, there was no effect of Rab7 on the Wnt inhibition mediated increase in EGFR recycling, suggesting that this change is not driven by diminished trafficking to late endosomes.

Enhanced endosomal recycling to the plasma membrane can increase the total and cell surface abundance of multiple receptor tyrosine kinases (44). To test if Wnts regulate the recycling of multiple RTKs, we used flow cytometry and indirect immunofluorescence staining of non-permeabilized HPAF-II cells. Indeed, we observed an increase in EPHA2, EPHB2, and EPHB4 on the surface of cultured HPAF-II cells (Fig 5E, S6L-M) after Wnt inhibition. The increase in the ephrin receptors at the cell surface is biologically relevant given their known role in providing positional cues for differentiation (50).

Next, we examined HPAF-II tumor lysates from mice treated with ETC-159 and observed an increase in the phosphorylation of EGFR at Y1068 and EPHA2 at Y588. This confirms that Wnt inhibition leads to the activation of EGFR and EPHA2 *in vivo* (Fig 5F-G). However, we were unable to measure any changes in the phosphorylation of other Ephs receptors due to the unavailability of reliable antibodies.

As a consequence of increased recycling, the total protein abundance of multiple RTKs was also increased following Wnt inhibition in HPAF-II orthotopic xenografts (Fig 5G-H). We observed an increase in the protein abundance of EGFR, ERBB2, EPHA2, EPHB4, and EPHB2, while many of these genes had no change in transcript abundance (Fig5G-H, S6N-O). This increase in the protein abundance of RTKs following Wnt inhibition was observed in multiple models, as there was also an increase in the protein abundance of ERBB2, EGFR, and multiple EPH receptors in PAXF1861 pancreatic patient-derived subcutaneous xenografts with an RNF43 G371fs mutation (Fig 5I). Notably, unlike what is observed in the xenografts, despite the increase in cell surface RTKs, there was minimal activation of phosphotyrosine following WNT inhibition in cultured cells. Taken together, our data indicate that Wnt signaling can regulate the recycling of multiple RTKs from late endosomes to the plasma membrane. The RTKs can then be activated by ligands present only in the local tumor environment.

### FAXDC2 regulates Wnt inhibition-mediated changes in RTK abundance

The cholesterol biosynthesis enzyme MSMO1, a C4-methyl sterol oxidase in the Bloch pathway, has previously been shown to regulate EGFR signaling in breast cancer cells (23, 51). To test if the RTK activation was dependent on changes in sterol metabolism in Wnt high cells, we examined the effect of the CYP51A1 inhibitor ketoconazole. Ketoconazole leads to the depletion of cholesterol biosynthesis intermediates downstream of lanosterol. Similar to what was observed with ETC-159 treatment, ketoconazole increased cell surface levels of EGFR in HPAF-II cells (Fig S7A), suggesting the importance of sterols in regulating RTK signaling in these cells. FAXDC2 changes the abundance of C4-methyl sterols, and FAXDC2-dependent genes were significantly enriched for endocytosis and vesicle-mediated transport processes (Fig 4D). We, therefore, tested with three approaches if FAXDC2 regulates RTK signaling in Wnt-addicted pancreatic cancer models.

First, we used cells with stabilized β-catenin that have low levels of FAXDC2 and high levels of lophenol and dihydro-T-MAS even upon ETC-159 treatment (Fig 1E and 2K-L). If FAXDC2 regulates RTK activation, we predicted that stabilized β-catenin would prevent the ETC-159 mediated increase in cell surface levels of RTKs. Indeed, stabilized β-catenin abrogated the increase in cell surface levels of multiple RTKs, including EGFR, EPHA2, and ERBB2, in HPAF-II cells and xenografts treated with ETC-159 (Fig 6A and S7B).

**Figure 6:**
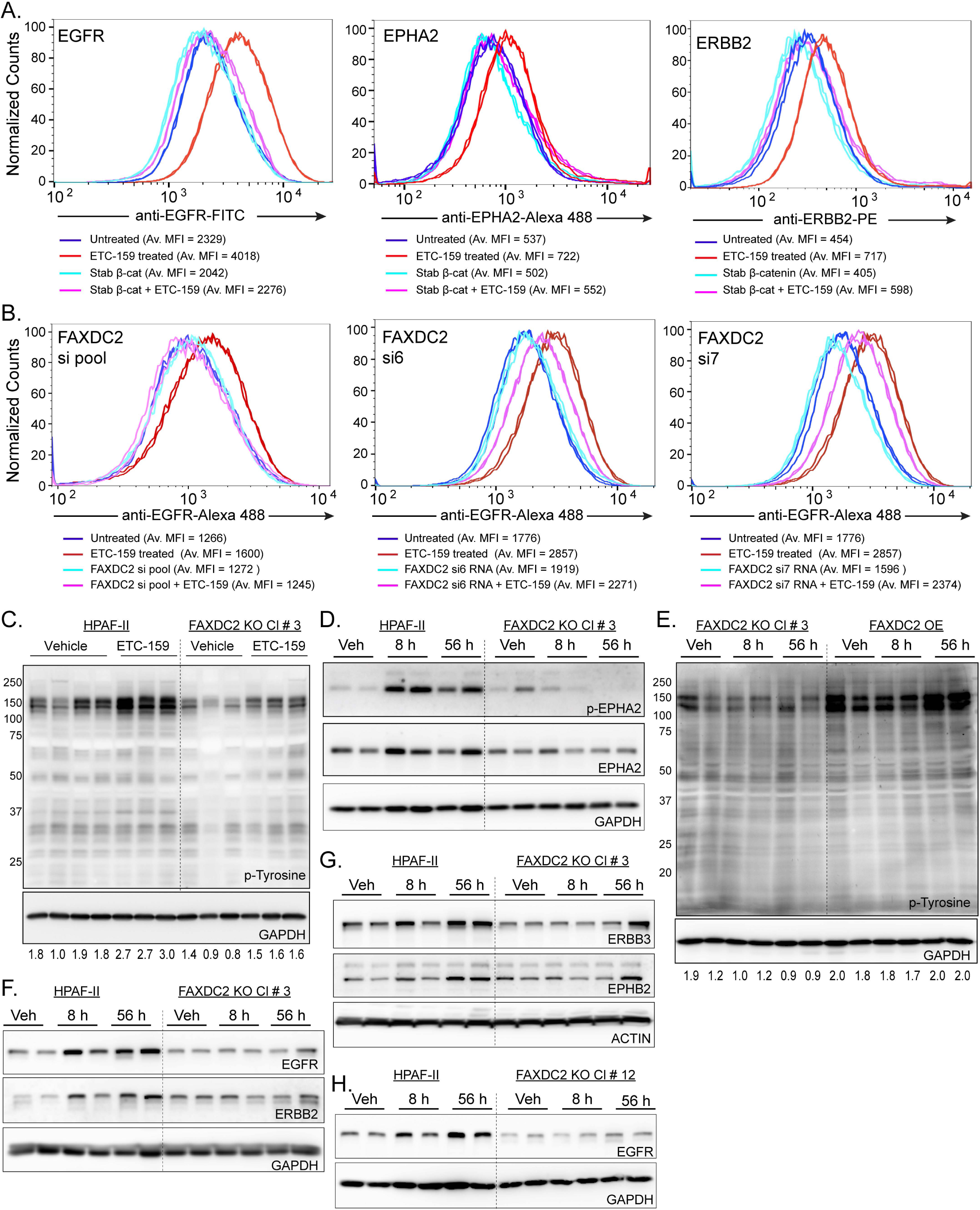
*FAXDC2* is downstream of Wnt/β-catenin in the regulation of RTK signaling. **A. *Stabilized β-catenin prevents multiple RTKs from moving to the cell surface upon treatment with ETC-159.*** The cell surface abundance of EGFR, ERBB2, and EPHA2 was assessed as before in HPAF-II cells with or without stabilized β-catenin. Cells were treated with DMSO or 100 nM ETC-159 for 72 h before staining with fluorescent tagged antibodies. Each histogram represents ∼50,000 cells. Data is representative of three independent experiments and two replicates are shown. **B. *Knockdown of FAXDC2 prevents Wnt inhibition-mediated increase in EGFR cell surface abundance***. HPAF-II cells were transfected with a pool of four siRNAs or two independent siRNAs (si6 and si7) against *FAXDC2* for 24 h, followed by treatment with ETC-159 for 72 h. The cells were then stained with Alexa Fluor 488-conjugated anti-EGFR antibody and analyzed by flow cytometry. Data is representative of three independent experiments, and two replicates are shown. **C-D. *Knockout of FAXDC2 in HPAF-II tumors prevents Wnt inhibition-mediated increase in tyrosine phosphorylation and phospho-EphA2 levels.*** Protein lysates from HPAF-II or FAXDC2 KO tumor xenografts from vehicle or ETC-159 treated mice were separated on 10% SDS gels and transferred to a PVDF membrane. The membranes were probed with anti-phosphotyrosine or phospho-EphA2 antibodies. Each lane represents lysate from an individual tumor. **E. *Overexpression of FAXDC2 in FAXDC2 KO tumors rescues the Wnt inhibition-mediated increase in tyrosine phosphorylation.*** Protein lysates from FAXDC2 KO or FAXDC2 OE tumor xenografts from vehicle or ETC-159 treated mice were probed with phosphotyrosine antibodies. Each lane represents lysate from an individual tumor. **F-H. *FAXDC2 KO prevents the Wnt inhibition-mediated increase in EGFR and Eph family receptor tyrosine kinase abundance.*** Protein lysates from HPAF-II or FAXDC2 KO tumors from mice treated with vehicle or ETC-159 were analyzed as in C. Data are shown from two independent FAXDC2 KO clones, (F-G) clone 3 and (H) clone 12. Each lane represents lysate from an individual tumor.

Second, we tested the direct effect of *FAXDC2* knockdown on EGFR levels *in vitro*. HPAF-II cells were transfected with a pool of siRNAs or two independent guide RNAs targeting *FAXDC2* in the presence or absence of ETC-159. As expected, Wnt inhibition increased cell surface EGFR levels, but the knockdown of *FAXDC2* prevented this increase in EGFR (Figs 6B and S7C-D). Notably, we observed that siRNA knockdown of *MSMO1* did not affect ETC-159-mediated changes in EGFR cell surface levels in HPAF-II cells (Fig S7E-F), suggesting that the observed effect is not a consequence of changes in overall cholesterol levels. This implies that FAXDC2 regulates cell surface receptor levels following Wnt inhibition in Wnt-ligand addicted models through its effect on KR-specific methyl sterols.

Third, we tested if FAXDC2 is required for the Wnt inhibition-mediated increase in RTK signaling in vivo. As before, there was a marked increase in tyrosine phosphorylation upon Wnt inhibition in HPAF-II tumors (Fig. 6C). Notably, in the absence of FAXDC2, the basal level of tyrosine phosphorylation was decreased, and the Wnt inhibition-mediated increase in tyrosine phosphorylation was also prevented (Fig 6C). We confirmed this result by examining EPHA2 tyrosine phosphorylation. EPHA2 was phosphorylated upon Wnt inhibition in parental tumors, but this increase in EPHA2 tyrosine phosphorylation was abrogated in the FAXDC2 KO tumors (Fig 6D). This effect of FAXDC2 knockout on RTK activation was so unexpected that to rule out off-target effects, we compared the phosphotyrosine levels in the FAXDC2 OE xenografts (FAXDC2 KO cells with FAXDC2 ectopically expressed under the control of CMV promoter). Consistent with a key role for FAXDC2 in regulating tyrosine phosphorylation, the FAXDC2 OE xenografts had enhanced tyrosine phosphorylation at basal levels, and there was no further increase by Wnt-inhibition (Fig 6E). Thus, changing FAXDC2 activity changes RTK activity.

Consistent with FAXDC2 being upstream of RTK recycling and signaling, FAXDC2 KO blocked the effect of Wnt inhibition on the protein abundance of multiple RTKs in the tumors, including EGFR, ERBB2, ERBB3, EPHA2, and EPHB2 (Fig 6D and F-G). Similar to clone #3, the increase in EGFR abundance after Wnt inhibition was also impaired in an independent FAXDC2 KO clone #12 tumors (Fig 6H).

These data are consistent with the model that Wnt/β-catenin signaling inhibits the expression of FAXDC2, and FAXDC2 controls the abundance of specific saturated C4-mono and dimethyl sterols to drive multiple RTKs to the plasma membrane where they are available for ligand interaction and activation.

### FAXDC2 is necessary and sufficient for Wnt inhibition-mediated MAPK activation

RTK phosphorylation leads to the activation of downstream signaling cascades, including Ras/ERK/MAPK signaling. Since knockout of FAXDC2 prevented the Wnt inhibition-mediated activation of multiple cellular tyrosine kinases, we tested how it impacted Wnt-regulated MAPK activation. We compared the phospho-ERK levels in control and FAXDC2-knockout tumors. As we previously reported, there was a significant increase in pERK1/2 levels upon Wnt inhibition in control tumors (Fig 7A-B) (16). FAXDC2 knockout tumors from both clones #3 and #12 had lower basal phospho-ERK than parental HPAF-II tumors (compare vehicle lanes in both groups, Fig 7A-B). Notably, the increase in ERK phosphorylation following Wnt inhibition was nearly abrogated in the FAXDC2 KO tumors compared to the WT HPAF-II tumors (Fig 7A-B). This was observed in two independent FAXDC2 knockouts, so it is neither a sgRNA nor clonal effect. The role of FAXDC2 in MAPK activation is consistent with its effect on RTK signaling, as FAXDC2 KO prevented the activation of RTKs. Thus, FAXDC2 expression is required for Wnt inhibition to activate both RTKs and MAPK.

**Figure 7:**
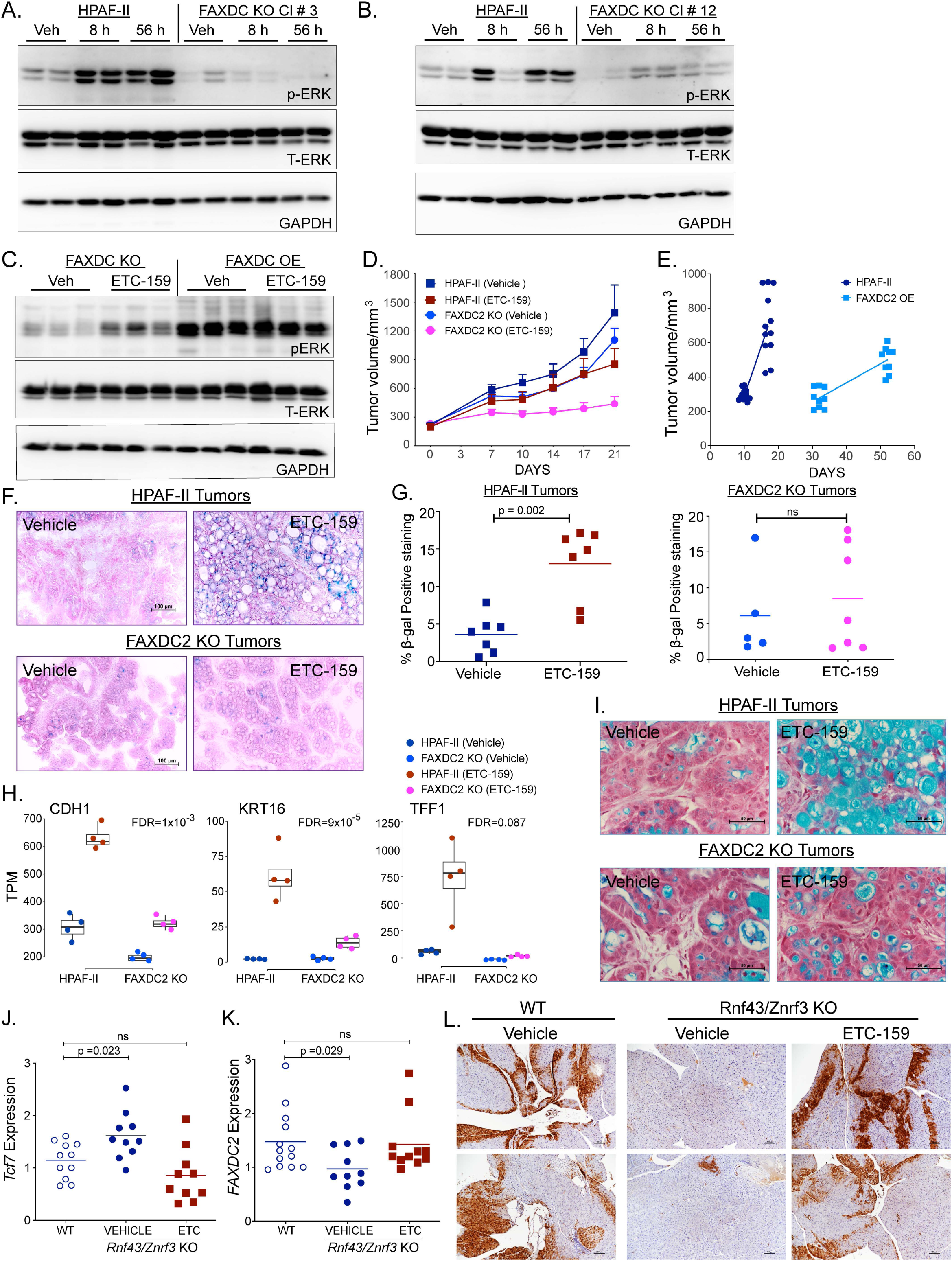
*FAXDC2* knockout prevents Wnt inhibition-induced MAPK activation, cellular differentiation, and senescence. **A-B. *FAXDC2 is required for MAPK activation.*** Xenografts from *FAXDC2* KO clones were generated using two independent guide RNAs. (A) Clone #3 and (B) clone #12 both had reduced ERK phosphorylation with no further increase upon ETC-159 treatment. Protein lysates from HPAF-II or FAXDC2 KO tumors from mice treated with vehicle or ETC-159 were analyzed by western blot. Each lane represents tumor lysate from an individual mouse. The protein lysates were prepared as a master mix and loaded on independent gels. Only one blot was probed for the load control. Load controls of Fig 7A is shared with Fig 6D and of Fig 7B is shared with Fig 6H. **C. *FAXDC2 overexpression mediates sustained ERK activation.*** Western blot analysis of protein lysates from HPAF-II or FAXDC2 OE tumor xenografts from mice treated with vehicle or ETC-159. Each lane represents tumor lysate from an individual mouse. **D. *Knockout of FAXDC2 reduced the growth of HPAF-II xenografts, with further reduction upon Wnt inhibition.*** HPAF-II or FAXDC2 KO cells were injected into the flanks of mice. Mice with established tumors were treated with vehicle or ETC-159 for 21 days. Volumes were recorded twice weekly, n= 5-6 mice/group. **E. *Overexpression of FAXDC2 delayed implantation and reduced the growth of HPAF-II xenografts.*** HPAF-II or FAXDC2 OE cells were injected into the flanks of mice. Tumor volumes were recorded after the tumors were established. n= 5-6 mice/group **F-G. *FAXDC2 knockout blunts the senescence response***. OCT-embedded sections of HPAF-II xenografts from mice treated with vehicle or ETC-159 for 21 days were stained for SA-β-galactosidase and counterstained with neutral red. Wnt inhibition caused an increase in SAβ-galactosidase in parental HPAF-II tumors; however, this was diminished in the FAXDC2 knockout tumors. (F) Representative images from the sections of each of the groups. (G) Tumor sections were scanned, and the percentage of the positively stained area (blue) was quantitated. Each dot represents the quantitation of a tumor section from an individual mouse, n=5-7 tumors/group. P values were calculated using the Mann-Whitney U test. **H-I*. FAXDC2 knockout blunts the differentiation response to Wnt inhibition.*** (H) Expression of selected differentiation markers was analyzed in tumors of all four groups by RNA sequencing. Each data point represents an individual tumor. FAXDC2 KO diminished the transcriptional response to Wnt inhibition. (I) FAXDC2 KO blunts the Wnt inhibition mediated increase in mucin expression. Representative images from the Alcian blue stained sections. **J-L. The Wnt-MAPK axis is present in the non-malignant mouse pancreas.** ***J-K. Genetic activation and subsequent pharmacologic inhibition of Wnt signaling in mouse pancreas leads to reciprocal regulation of Faxdc2 expression*.** Pancreas from control or ETC-159 treated WT and *Ptf1a^Cre^Rnf43*^-/-^*Znrf3^-/-^*mice were analysed for *Tcf7* and *Faxdc2 (Gm12248) expression* by qRT-PCR. *Knockout of Rnf43 and Znrf3 in mouse pancreas* led to an increase in the expression of Wnt target gene *Tcf7* (K) and reduction in *Faxdc2* (L). Wnt inhibition with ETC-159 treatment rescues the altered expression of both genes in the pancreas of *Ptf1a^Cre^Rnf43*^-/-^*Znrf3^-/-^* and restores it to levels comparable to the WT mice pancreas. Data is from two independent biological experiments with each dot representing an individual mouse, n=5-6 mice/group. **L**. ***Wnt activation leads to reduction in p-ERK levels in the mouse pancreas.*** MAPK activity was assessed by IHC with anti-p-ERK antibody. Representative image p-ERK stained pancreas from WT*, Ptf1a-Cre-Rnf43/ Znrf3* KO mice and ETC-159 treated *Ptf1a-Cre-Rnf43/ Znrf3* KO mice. MAPK activity in the pancreas is suppressed by genetic activation of Wnt signaling in *Ptf1a-Cre Rnf43/Znrf3* knockout mouse pancreas and restored upon Wnt-inhibition with ETC-159 treatment to levels comparable to WT mice.

To test if ERK activation was a direct consequence of FAXDC2 expression independent of Wnt signaling, we compared the p-ERK levels in FAXDC2 OE xenografts, where both lophenol and T-MAS levels are markedly suppressed (Fig 3G-H). Overexpression of FAXDC2 led to high basal p-ERK levels (Fig 7C), and there was no further increase upon Wnt inhibition, consistent with the effect on tyrosine phosphorylation (Fig 6E),

Taken together with the enrichment of ETS and AP1 binding sites in the genes regulated by FAXDC2, these results show that the regulation of FAXDC2 expression by the Wnt/β-catenin axis is required for maintaining endocytic recycling and activation of RTKs and downstream MAPK signaling.

### The role of FAXDC2 in Wnt-inhibition-mediated senescence and differentiation

RTK/RAS/MAPK signaling can drive proliferation, but this pathway is also essential for differentiation in multiple settings. RAS signaling in cancer is modulated for optimal growth and has a “sweet spot;” loss of activity prevents proliferation but excessive activity can cause senescence (52–54). Consistent with decreased basal p-ERK signaling, FAXDC2 KO cells grew ∼30% slower than the parental cells both in culture and as tumors in the flanks of immunocompromised mice (Fig 7D). Conversely, FAXDC2 overexpressing cells with markedly increased phosphotyrosine and phospho-ERK also grew very slowly in mice. The FAXDC2 OE cell pool took 5-6 weeks to establish tumors in the mice, unlike the parental HPAF-II cells that formed tumors in 5-6 days (Fig 7E). The FAXDC2 OE tumors increased in volume by 1.9-fold in three weeks compared to the parental HPAF-II tumors, which more than doubled (2.3-fold) in one week.

We and others have previously demonstrated that Wnt inhibition causes both differentiation and cellular senescence in Wnt-addicted cancers (14, 15, 55). Here, we tested the role of FAXDC2. As before, treatment of HPAF-II tumors with Wnt inhibitor significantly increased senescence-associated β-galactosidase (SA-β-gal) levels as well as the expression of cytokines and growth factors implicated in senescence-associated inflammation (Fig 7F-G, S8A). Knockout of FAXDC2 prevented this Wnt-inhibition-driven senescence. Several secreted growth factors and cytokines, including *IL32, HBEGF, ILK*, and *CTGF* implicated in senescence-associated inflammation were no longer upregulated upon Wnt inhibition in FAXDC2 knockout tumors (Fig S8A) (56–59). This suggests that the FAXDC2-RTK-MAPK pathway regulates Wnt inhibition-mediated senescence.

FAXDC2 appears to be similarly required for the differentiation seen after Wnt withdrawal. Inhibition of Wnt signaling leads to the differentiation of Wnt-ligand-addicted tumors, demonstrated in part by enhanced staining for mucins and increased expression of differentiation markers such as *CDH1, CDHR2, TFF1* and *KRT16* (Fig 7H-I, S8B). Notably, in the absence of FAXDC2, there was markedly diminished upregulation of these markers. Consistent with this, there was no increase in the staining for mucins in the ETC-159 treated FAXDC2 KO tumors. Thus, although tumor proliferation was partially affected by FAXDC2 knockout, the differentiation and senescence response required the increased expression of FAXDC2.

Taken together, these data show that FAXDC2 is a Wnt-repressed enzyme that regulates the abundance of specific C4-methyl sterols to facilitate cellular senescence and differentiation via hyperactivation of the RTK-MAPK signaling pathway.

### FAXDC2 regulates MAPK signaling in Wnt-driven hyperplasia of the pancreas

We asked if the Wnt/FAXDC2/MAPK pathway was also present in non-malignant tissues. To do this, we examined a mouse genetic model of activated Wnt-dependent signaling in the pancreas. We generated mice with homozygous deletion of *Rnf43* and *Znrf3* in the pancreas (*Ptf1a*^Cre^*Rnf43*^fl/fl^/*Znrf3*^fl/fl^) (60). These mice developed modest pancreatic hyperplasia (Fig S8C) but not pancreatic cancer. As expected, they also showed an enhanced response to endogenous Wnts as evidenced by increased expression of the Wnt target gene *Tcf7* (Fig 7J). Consistent with *Faxdc2* being a Wnt-repressed gene in non-malignant tissues, there was a ∼33% decrease in *Faxdc2* (*Gm12248*) *expression* in the *Ptf1a*^Cre^*Rnf43*^fl/fl^/*Znrf3*^fl/fl^ mice (Fig 7K). Furthermore, the increase in Wnt signaling as a result of *Rnf43*/*Znrf3* deletion was accompanied by a loss of basal phospho-ERK staining, most prominent at the periphery of the pancreatic lobules (Fig 7L). Blocking Wnt secretion by 21 days of treatment with ETC-159 in the *Ptf1a*^Cre^*Rnf43*^fl/fl^/*Znrf3*^fl/fl^ mice restored *Tcf7* and *Faxdc2* expression to normal levels. Importantly, this ETC-159 mediated restoration of *Faxdc2* expression was accompanied by recovery of p-ERK staining. Thus, Wnt signaling also suppresses FAXDC2 and MAPK signaling in non-malignant tissues, indicating that modulation of cholesterol biosynthesis intermediates may be a Wnt effector pathway with widespread importance.

## DISCUSSION

Wnt signaling can regulate the balance between proliferation and differentiation in stem cells and cancers. Here, taking advantage of a potent Wnt inhibitor and a comprehensive transcriptomic analysis of a Wnt-ligand-addicted orthotopic cancer model, we identified a core Wnt/β-catenin repressed cholesterol biosynthesis enzyme, the 4-methyl sterol monooxidase FAXDC2, that controls this balance. Mechanistically, we find that inhibition of Wnt secretion led to increased abundance and activity of receptor tyrosine kinases including EGFR and EPHA2 at the cell surface and enhanced signaling via the RAS-MAPK cascade to drive widespread increases in gene expression. Upstream of these steps, we find that loss of *FAXDC2* prevents Wnt-regulated RTK activation and downstream MAPK signaling to prevent a differentiation/senescence response.

In multiple cancers, oncogenic mutations of RAS lock it in the GTP-bound ON state, leading to activated MAPK signaling. Recent studies have demonstrated that RAS signaling is tightly regulated even in cancer, in part because excessive RAS signaling drives senescence (52–54). One unexpected finding of our recent study was that high Wnt signaling in cancers prevented hyperactivation of MAPK signaling and Ras-mediated senescence. In this context, our data suggest that one mechanism for RAS-mutant cancers to avoid excess signaling and senescence is to concurrently repress FAXDC2, thereby preventing RTK hyperactivation and inhibiting downstream MAPK signaling.

FAXDC2 is a paralog of MSMO1/SC4MOL in cholesterol biosynthesis and is required for the demethylation of the C4 position of cholesterol biosynthesis intermediates (Fig 2) (18). Not much is known about the function of FAXDC2 apart from a single report implicating it in megakaryocyte differentiation (61). FAXDC2 is both ubiquitously expressed and highly conserved through evolution in plants, animals, and sponges and contains the signature sequences present in sterol monooxidases (Fig S2). We find that *FAXDC2* expression regulates the abundance of specific C4-methyl sterols, lophenol, and dihydro-TMAS. These specific sterols are saturated at the 24,25 position, placing them in the Kandutsch and Russell branch of cholesterol biogenesis (19, 20). Hence, FAXDC2 may preferentially act on the cholesterol precursors in the Kandutsch-Russell arm of the cholesterol biosynthesis pathway. We note that Kandutsch and Russell initially identified 24,25 dihydro-cholesterol precursors in cancer tissue, calling them tumor sterols (19, 20). Consistent with this, we find that lophenol and dihydro-TMAS are elevated in pancreatic and colorectal cancers (Figs 2 and 3).

Lophenol is widely present in plants such as aloe vera that are ascribed medicinal properties, including activity in metabolic disorders (e.g., see (62)). In *Caenorhabditis elegans*, lophenol is required for dauer formation. Our data suggest a potential biological mechanism for the effects of these C4-methyl sterols in RTK/MAPK regulation, a subject we are actively investigating.

FAXDC2 has not previously been identified as a Wnt repressed gene. We were able to identify it as a Wnt/β-catenin target due to several factors. First, our Wnt-regulated dataset was obtained from *in vivo*, orthotopic Wnt-addicted cancer models and patient derived xenografts, where the dynamic range of Wnt-regulated genes was much higher than that seen in subcutaneous xenografts and cultured cells (14). Second, our discovery dataset contained 42 independent orthotopic tumors sequenced to high depth, giving us robust statistical power. We validated our findings in independent datasets that identified Wnt target genes using genetic rather than pharmacological approaches. Finally, we confirmed the observational findings in independent experiments using stabilized β-catenin. Hence, our identification of FAXDC2 as a Wnt/β-catenin repressed gene in pancreatic and colorectal cancers is made with high confidence. Determining if FAXDC2 is a direct or indirect β-catenin target gene is an area of active investigation. This is not as straightforward as it appears, as important β-catenin/TCF4 binding sites are now well-recognized to be in enhancers that are quite distant from transcription start sites (63, 64).

We and others have reported that Wnt inhibition drives differentiation of Wnt-addicted cancers. Reactivation of Ephrin-Eph bidirectional signaling following Wnt inhibition could be one potential mechanism contributing to the differentiation of tumors. EPH signaling is well known to provide critical positional cues to cells during development, migration, and differentiation, including proliferation in the intestinal stem cell niche (65, 66). Future studies will test if the FAXDC2-dependent increase in abundance of Ephs on the cell surface and consequent engagement of bidirectional signaling plays a crucial role in the differentiation response following Wnt inhibition.

MYC is a well-recognized and essential target of Wnt signaling. We previously identified 2,131 genes regulating cell cycle and ribosomal biogenesis whose transcriptional response to Wnt inhibition in vivo depended on MYC status (14). Our study shows that de-repression of FAXDC2 is also a key consequence of Wnt inhibition, suggesting FAXDC2 may be as important a Wnt target as MYC in Wnt-high cancers. Confirming the importance of the Wnt/FAXDC2 axis in regulating the senescence/differentiation response, knockout of FAXDC2 alters the expression of ∼3,500 Wnt-regulated genes. Thus, activation of MYC and repression of FAXDC2 may be central to Wnt signaling for regulating the balance between proliferation vs. senescence and differentiation.

Our study provides the first evidence for Wnt-regulated repression of the expression of FAXDC2, leading to an altered abundance of C4 methyl sterols and RTK/MAPK signaling. Future studies will address the importance of FAXDC2 and the C4-methyl sterols it regulates during development and adult tissue homeostasis.

## METHODS

### Flow Cytometry

For flow cytometric analysis, cells were seeded in 60 mm dishes and transfected with 50 uM siRNAs against FAXDC2, MSMO1, RAB11A, RAB11B, RAB4, RAB25 or EHD1 (Qiagen). 24 h after transfection, the media was replaced with fresh media containing DMSO or 100 nM ETC-159. After 72 h of treatment, non-permeabilized cells were incubated with anti-EGFR-FITC (SC-120, Santa Cruz Biotechnology, Dallas, TX), ERBB2-PE (98710, Cell Signaling Technology, Danvers, MA), EphA2 (ab150304, Abcam, Cambridge, UK) or EphB2-APC (564699, BD Biosciences, San Jose, CA) antibodies for 60 min. Cells were washed with PBS followed by incubation with goat anti-rabbit IgG (H+L) Alexa fluor-594 secondary antibody (A-11012 from Thermo Scientific (Rockford, IL) was used for EphA2. Samples were acquired on a BD Fortessa, and the data were analyzed using FlowJo software v10.0.7.

### Co-localization assays

Epitope-tagged FAXDC2, MSMO1, and NSDHL cDNA were cloned into a pcDNA3.1 mammalian expression vector using restriction enzymes. All tags were located at the N-terminus. FAXDC2 and MSMO1 were tagged with V5. NSDHL was tagged with Flag. 75,000 HeLa cells were plated on coverslips in 12-well plates and transiently transfected with the relevant plasmids (MSMO1 and NSDHL or FAXDC2 and NSDHL) using lipofectamine 2000 (10 ng of plasmid per well). Forty-eight hours after transfection, cells were fixed in 4% paraformaldehyde and permeabilized using a commercially available saponin-based permeabilization and blocking reagent (1x BD Perm/Wash buffer #554723). Cells were then stained using primary antibodies against Flag (Proteintech 20543-1-AP; 1:2000) and V5 (BioRad MCA1360; 1:500) and subsequently with fluorophore-labeled anti-mouse (goat anti-mouse Alexa Fluor 488; 1:2000; Invitrogen) and anti-rabbit secondary antibodies (goat anti-rabbit Alexa Fluor 594; 1:2000; Invitrogen). Duolink Mounting Medium with DAPI was used to mount the coverslips. Images were obtained using a Nikon Ti2-E microscope with a 100x oil immersion objective and were analyzed using NIS-Elements software.

### Generation of MSMO and FAXDC2 KO cells

To generate FAXDC2 or MSMO KO HPAF-II cells, MSMO1 sg RNA (sg: 5’-CACCGCAGAGACATGGGAAAACCAA-3’) or FAXDC2 sg RNAs (sg3 5’-TCTTGTTCTACTATTCACAC-3’, sg4-TGGGGAAAGATATCATGCAC-3’) were cloned into was cloned into FgH1tUTG or FgH1tUTCyan plasmid (Addgene #85551) plasmids respectively. HPAF-II cells stably expressing FUCas9Cherry (Addgene#70182) were transduced with MSMO1 sg RNA or FAXDC2 sgRNA along with pMD2.G (Addgene #12259) and psPAX2 (Addgene #12260) plasmids. The transduced cells were sorted using flow cytometry to select GFP or Cyan-positive cells. Genome editing was validated by PCR amplification and sequencing.

### Low-Density Assays

9000 cells/well were plated in 24-well cell culture plates for low-density assays. HPAF-II cell pools with inducible single guide (isg) RNAs against FAXDC2 or MSMO1 or FAXDC2 KO cells with isgRNAs against MSMO1 (FAXDC KO isgMSMO) were plated in growth media with delipidated FBS (Gemini Bio 900123). The cells were treated with 2.5 μg/ml doxycycline (DOX) or DMSO and were allowed to grow for ∼10 days. The cells were then stained with 0.5% crystal violet solution (Sigma-Aldrich, V5265), and the images were acquired on GelCount (Oxford Optronix, Abingdon, UK). Crystal violet was solubilized by incubating with 0.1 % SDS for 4-5 hrs at RT, and the absorbance was measured at 570 nM using BioRad XMarkTM Microplate Spectrophotometer.

### Cholesterol Assay

To measure total cholesterol, 400,00 cells/well were plated in 6-well cell plates in growth media with delipidated FBS and harvested after treatment with either 2ug/ml of doxycycline or DMSO. To extract lipids, 1 million cells were resuspended in 200ul of Chloroform: Isopropanol: NP-40 (7:11:0.1, v/v) followed by centrifugation at 15,000 g for 10 mins at room temperature. After solvent extraction, the samples were dried in a Speedvac at 50°C for 30 mins to remove residual traces of organic solvent. The dried lipid extracts were dissolved in 200 ul assay diluent, and total cholesterol was measured using Amplex® Red Cholesterol Assay Kit (Invitrogen, Catalog No. A12216) per the manufacturer’s protocol.

### Data analysis

Data were analyzed using Prism v5.0 (GraphPad, La Jolla, CA, USA) and R/Bioconductor. Significance for all tests was set at p ≤ 0.05 unless otherwise stated. *p ≤ 0.05, **p ≤ 0.01,***p ≤ 0.001,****p ≤ 0.0001 in all instances. Biorender was used to draw the graphical abstract.

### Study Approvals

The Duke-NUS Institutional Animal Care and Use Committee approved the animal studies and complied with applicable regulations. Institutional review boards of Singhealth (2018–2795) approved the analysis of CRC samples.

## ACKNOWLEDGMENTS

We gratefully acknowledge the assistance, advice, and support of the Petretto and Virshup lab members. Wenxu Zhou in the Nes lab performed the initial sterol analysis. We gratefully acknowledge the technical assistance of Shreya Sridharan with the biological assays and mouse studies, the Duke-NUS vivarium staff, and the exceptional contributions of the DUKE-NUS metabolomic facility. ETC-159 was a generous gift from the Experimental Drug Development Centre, A*STAR, Singapore. We thank Andreas Hecht, Albert-Ludwigs University Freiburg, Germany, for the generous gift of TCF7L2 knockout, and control HCT116 and HT29 cell lines and Bon-Kyoung Koo, Vienna Biocenter, Austria, for the Rnf43 ^fl/fl^*/Znrf3*^fl/fl^ mice.

This research is partly supported by STAR Award MOH-000155 to DMV from the National Research Foundation Singapore, administered by the Singapore Ministry of Health’s National Medical Research Council. BM acknowledges the support of the Singapore Ministry of Health’s National Medical Research Council Open Fund–Independent Research grant NMRC/OFIRG/0055/2017. David Nes acknowledges the support of WDN-R21(33)AI119782. Iain B Tan is supported by National Medical Research Council MOH-000425-00 (CTG-ICT) and the National Cancer Centre Research Fund.

## Data availability

The RNA-Seq data were deposited in the NCBI’s Sequence Read Archive under accession number PRJNA549884. RNAseq analysis code is available at: https://github.com/harmstonlab/wnt_faxdc2_manuscript

## Author contributions

BM and DMV designed the study. BM, SW, and SP performed animal studies and biochemical assays. NH performed the bioinformatics analysis. WDN performed sterol structure identification and bioinformatics of FAXDC2 versus MSMSO1/SMO1 and contributed extensively to biosynthesis reasoning. ET and IBHT provided the human CRC tissue samples. EP, BM, and DMV supervised the study. BM, NH, and DMV wrote the manuscript with significant input from WDN. All authors read and approved the manuscript.

