## Supplemental data for "The cholesterol biosynthesis enzyme FAXDC2 couples Wnt/β-catenin to RTK/MAPK signaling"

#### SUPPLEMENTAL METHODS

##### Tissue Culture

HEK293 cells were obtained from ATCC and grown in DMEM, EMEM, or RPMI-1640 supplemented with 10% fetal bovine serum and 1% GlutaMAX (Life Technologies, Grand Island, NY) in a 37°C humidified incubator with 5% CO2. AsPC-1 cells from ATCC were grown in RPMI supplemented with 10% fetal bovine serum in a 37°C humidified incubator with 5% CO2. HPAF-II cells from ATCC were grown in EMEM supplemented with 10% fetal bovine serum in a 37°C humidified incubator with 5% CO2. HCT116 TCF7L2KO, HT29 TCF7L2 KO, and their parental cell lines were generous gift from Dr. Andreas Hecht (1). E[beta]P (Addgene plasmid # 24313) was used for generating HPAF-II cells stably overexpressing stabilized β-catenin as described previously (2). All cell lines were regularly tested for mycoplasma contamination and confirmed mycoplasma free.

##### Animal care

NOD SCID gamma (NSG) mice were purchased from InVivos, Singapore, or Jackson Laboratories, Bar Harbor, Maine. The Duke-NUS Institutional Animal Care and Use Committee approved the animal studies and complied with applicable regulations. Animals were housed in standard cages and were allowed access ad libitum to food and water. Rnf43 fl/fl/Znrf3fl/fl mice were generously provided by Bon-Kyoung Koo and Hans Clevers (3), and Ptf1a Cre mice were obtained from Jackson Laboratories, Bar Harbor, Maine. Institutional review boards of Singhealth (2018-2795) approved the analysis of CRC samples.

##### Tumor growth and mice treatment

Mouse xenograft models were established by orthotopic injection of HPAF-II or AsPC-1 cells in NSG mice as described previously (4). For tumor growth studies, HPAF-II, FAXDC2 knockout, or FAXDC2 overexpressing (OE) HPAF-II cells were suspended in 50% matrigel and injected subcutaneously into the flanks of NSG mice. Mice were treated with ETC-159 after the establishment of tumors. ETC-159 was formulated in 50% PEG 400 (vol/vol) in water and administered by oral gavage at a dosing volume of 10 μL/g body weight as previously described (4). The tumor dimensions were measured with a caliper routinely, and the tumor volumes were calculated as 0.5 x length x width2 as described (5) previously. All mice were sacrificed 8 hours after the last dosing. At sacrifice, tumors were resected, weighed, and snapped frozen in liquid nitrogen or fixed in 10% neutral buffered formalin.

##### Western Blot analysis

For immunoblot analysis, tumors were homogenized in 4% SDS buffer using a polytron homogenizer. An equal amount of proteins was resolved on 10% SDS-polyacrylamide gel and transferred to PVDF membranes. Western blots were performed according to standard methods. The blots were probed with phospho-p44/42 MAPK (Erk1/2) (Thr202/Tyr204) antibody (9106), p44/42 MAPK (Erk1/2) antibody (9102), EphA2 (6997), pEhA2 (12677), pEGFR (48576) EphB2 (83029), EphB4 (14960), ErbB2 (4290), ErbB3 (12708), and Phospho-Tyrosine (8954) antibodies from Cell Signaling Technologies, Danvers, MA; EGFR antibody (1005) from Santa Cruz Biotechnology, Dallas, TX; actin (3280) and GAPDH (8245) antibodies from Abcam and FAXDC2 antibody (22046-1-AP ) from Proteintech. Blots were developed using SuperSignal West Femto or SuperSignal West Dura substrate (Thermo Scientific; Rockford, IL). Protein lysates from HPAF-II orthotopic xenografts were analysed using a phosphotyrosine antibody array (ab193662) from Abcam, Waltham, MA according to the manufacturer's instructions. Images were captured digitally using the LAS-3000 Life Science Imager (Fujifilm; Tokyo, Japan). "Restore western blot stripping buffer" from Thermo Fisher was used for stripping the blots for reprobing.

#### RNA Isolation and qRT-PCR

Tumors were homogenized in RLT buffer using a polytron homogenizer, and total RNA was isolated using an RNAeasy kit (Qiagen, Germany) according to the manufacturer's protocol. The RNA-seq libraries were prepared using the Illumina TruSeq stranded Total RNA protocol with subsequent PolyA enrichment. For qRT-PCR, RNA was reverse transcribed with iScript reverse transcriptase (BioRAD, Hercules, CA, USA). Real-time quantitative PCR (qPCR) was performed using SsoFast™ EvaGreen® assay Supermix from BioRad (Hercules, CA, USA). EPN1 and ACTB were used as housekeeping genes. The primers used are listed in Table S3.

#### Immunofluorescence

For immunofluorescence analysis, $4x10^5$ HPAF-II cells were plated in 24 well plates on glass coverslips. Cells were treated with DMSO or 100 nM ETC-159. After 72 h, the cells were fixed with 4% neutral buffered formalin (157-4, Electron Microscopy Sciences, PA) for 15 min at RT. The fixed cells were washed with PBS followed by blocking for 1 hour with 2% BSA/0.1% Triton X-100/PBS and then incubated with anti-EGFR-Alexa Fluor 488 antibody (1:200, SC-120, Santa Cruz Biotechnology, Dallas, TX), anti-EPHB2 (1:300 dil) or anti-EPHB4 (1:300 dil) primary antibodies overnight in 2% BSA/PBS at 4°C. After incubation with primary Ab, cells were stained with goat anti-Rabbit IgG (H+L) Alexa Fluor 594 (#A-11012, Thermo Scientific Rockford, IL) secondary antibody at room temperature for 1 h. To visualize, glass coverslips were mounted in ProLong Diamond Antifade Mountant with DAPI (Invitrogen #P36962) and then imaged with Leica (TCS SP8) inverted confocal microscope at 100X magnification.

#### Immunohistochemistry

Tissue sections of formalin-fixed paraffin-embedded tumors were deparaffinized in xylene and rehydrated in ethanol gradients. For p-ERK staining, after antigen retrieval with sodium citrate buffer pH 6.0 for 10 min, the endogenous peroxidase activity was blocked by incubation with H2O2. The sections were then incubated overnight with 1:200 diluted phospho-p44/42 MAPK antibody (4376S, Cell Signaling Technologies, Danvers, MA) followed by HRP conjugated anti-rabbit secondary antibody for one h. Incubation with 3,3'-diaminobenzidine chromogen substrate resulted in brown staining of phospho-ERK positive cells, and the nuclei were counterstained with Mayer's hematoxylin. Brightfield images were acquired on a Nikon Eclipse Ni-E microscope. OCT-embedded tissue sections were stained for SA-β-galactosidase using the protocol described previously (2).

#### Sterol extraction and GC/MS analysis for sterol identification

To extract the sterols, the tumors were saponified with 1 ml of 10% methanolic KOH at 80°C for two hours, followed by extraction with 1 ml n-hexane. One ug of epicoprostanol (0.1 mg/ml in toluene) was added as an internal standard. The sterols in the mixture were extracted with 1 ml of n-hexane and dried under nitrogen. The residues were stored in a CaCl2 desiccator for 24 h to remove moisture. The extracted sterols were then converted to their trimethyl silane esters using N, O-Bis (trimethylsilyl), trifluoroacetamide (BSTFA) with trimethylchlorosilane (TMCS) (99:1), and pyridine as a catalyst. For analysis, each derivatized sample (5 µl) was injected into the 0.25 mm capillary column of GC/MSD (Agilent GC 6890, fitted with an automatic Liquid Sampler and a 5973 quadrupole MS detector (Agilent Technologies). The inlet temperature was 250˚C, and the helium carrier gas flowed at a constant rate of 1.2 mL/min. The GC oven temperature was set at 170˚C initially for 1 min with an increase to 280˚C at a rate of 40˚C min-1 and then held for 20 min. The transfer line temperature was set at 280˚C, MS source at 230˚C, and the quadrupole temperature at 150˚C. Ionization was by electron impact at 70 eV. The mass calibrant perfluorotributylamine was used to auto-tune the MSD. The MSD was run at scan mode ranging from 50 to 550 amu.

The data was analyzed using Agilent GC/MSD Productivity Chemstation software and Automated Mass Spectral Deconvolution and Identification System (AMDIS). The sterols were identified by comparing their retention times and mass spectra to the authentic standards and commercial GC/MS database (NIST/EPA/NIH Mass Spectral Library, NIST 08). The sterols were identified by comparing their retention times and mass spectra to the authentic standards, commercial GC/MS database (NIST/EPA/NIH Mass Spectral Library, NIST 08), or purified 4,4-dimethylcholesta-8,24-dienol (isolated from the Saccharomyces cerevisiae ERG25 mutant (6), Figure S5E). The database has no sample for 4,4-dimethylcholest-8-enol, and no standard was available. However, the mass spectrum of this sterol was similar to 4,4-dimetylcholesta-8,24-dienol but two mass units higher, and it eluted about 1.5 min early, indicating that it has a saturated side chain. The strong m/z=135 was from the 4,4-dimethyl substitution. The mass spectrum of the targeted sterol showed high similarity to that of the published mass spectrum of 4,4-dimethylcholest-8-enol (7), and it was confirmed to be 4,4-dimethylcholest-8-enol.

#### Semi-quantitative analysis of sterols

The selective ion monitoring (SIM) method was developed to quantify the trace amount of sterols in the tumor tissues with high sensitivity (detection limit of 10 ng on column). The first group of ions was from 9 min to 14.6 min, and m/z 355 amu for the internal standard was monitored with 25% dwell time; the second group of ions was from 14.6 min to 28 min with m/z 458 amu for cholesterol and lathosterol, m/z 472 for campesterol and lophenol, m/z 484 for 4,4-dimethylcholesta-8,24-dienol and m/z 486 for 4,4-dimethylcholest-8-enol and sitosterol. Each ion was monitored with a 25% dwell time. The intensity of each ion was integrated using the Chemstation software, and TIC (total ion current), which represents the amount of each sterol, was calculated by the conversion factor of the SIM to TIC derived from the standards (the conversion factor for 4,4-dimethylcholesta-8,24-dienol was used for the calculation of 4,4-dimethylcholest-8-enol since their mass spectra were very similar). The amount of each sterol was normalized to the internal standard and the fresh tissue weight. The amount of cholesterol could not be accurately determined because of its high concentration in the tissue, and the mass detector was saturated even in SIM mode.

#### RNA-seq analysis

Data processing and QC. RNA-seq datasets were analyzed and clustered as described in Madan et al. (4). The RNA-seq of FAXDC2 KO tumors was performed using the same pipeline. Sequences were assessed for quality, and reads from mouse (mm10) were removed using Xenome (8). The remaining reads were aligned against hg38 (Ensembl version 79) using STAR v2.5.2 (9) and quantified using RSEM v1.2.31 (10). Reads mapping to chrM or annotated as rRNA, snoRNA, and snRNA were removed. Genes with less than ten reads mapping on an average were removed. Differential expression analysis was performed using DESeq2 (11). Independent filtering was not used in this analysis. Pairwise comparisons were performed using a Wald test. To control for false positives due to multiple comparisons in the genome-wide differential expression analysis, we used the false discovery rate (FDR) computed using the Benjamini-Hochberg procedure. Gene-level counts were transformed using a variance-stabilizing transformation and converted to z-scores. Time was transformed using a square root transformation. All genes differentially expressed over time (DESeq2, false discovery rate (FDR) < 10%) were clustered using GPClust (12) using the Matern32 kernel with a concentration (alpha) parameter of 0.001 and a length scale of 6.5.

Fold changes for all FAXDC2-dependent genes (interaction test, FDR < 10%) were clustered using hierarchical clustering with complete linkage and correlation distance. We used the average silhouette width to determine this dataset's optimal number of clusters (Figure S8, N=6 tumors/group).

#### Functional enrichment analysis

For the analysis of the Wnt-repressed genes (Figure 1), Gene Ontology (GO) enrichments were performed using GOStats (13) using all genes differentially expressed (FDR < 10%) as background. ReactomePA (14) was used for investigating pathway enrichments using the same background. Terms with an FDR < 10% were defined as significantly enriched. For the analysis of the FAXDC2-dependent genes (Figure 8), functional enrichments were performed using all expressed genes as background.

#### Transcription factor binding sites (TFBS) analysis

TFBS motifs were obtained from the JASPAR2018 database (15). Promoters were defined as 500 bp upstream and downstream from the ENSEMBL annotated transcription start site. AME was used to search for enriched motifs in these regions using all expressed genes not in a specific cluster as background and a hit-lo-fraction of 0.5. p-values reported by AME were corrected for multiple testing using FDR (16). Motifs with an FDR < 10% were defined as significantly enriched. A complete list of the motif enrichments for the Wnt-repressed clusters C2, C4, C6, and C8 is reported in Supplemental Table S4. A complete list of motif enrichments for clusters of genes identified as being FAXDC2-dependent is reported in Supplemental Table S2.

Figure S1

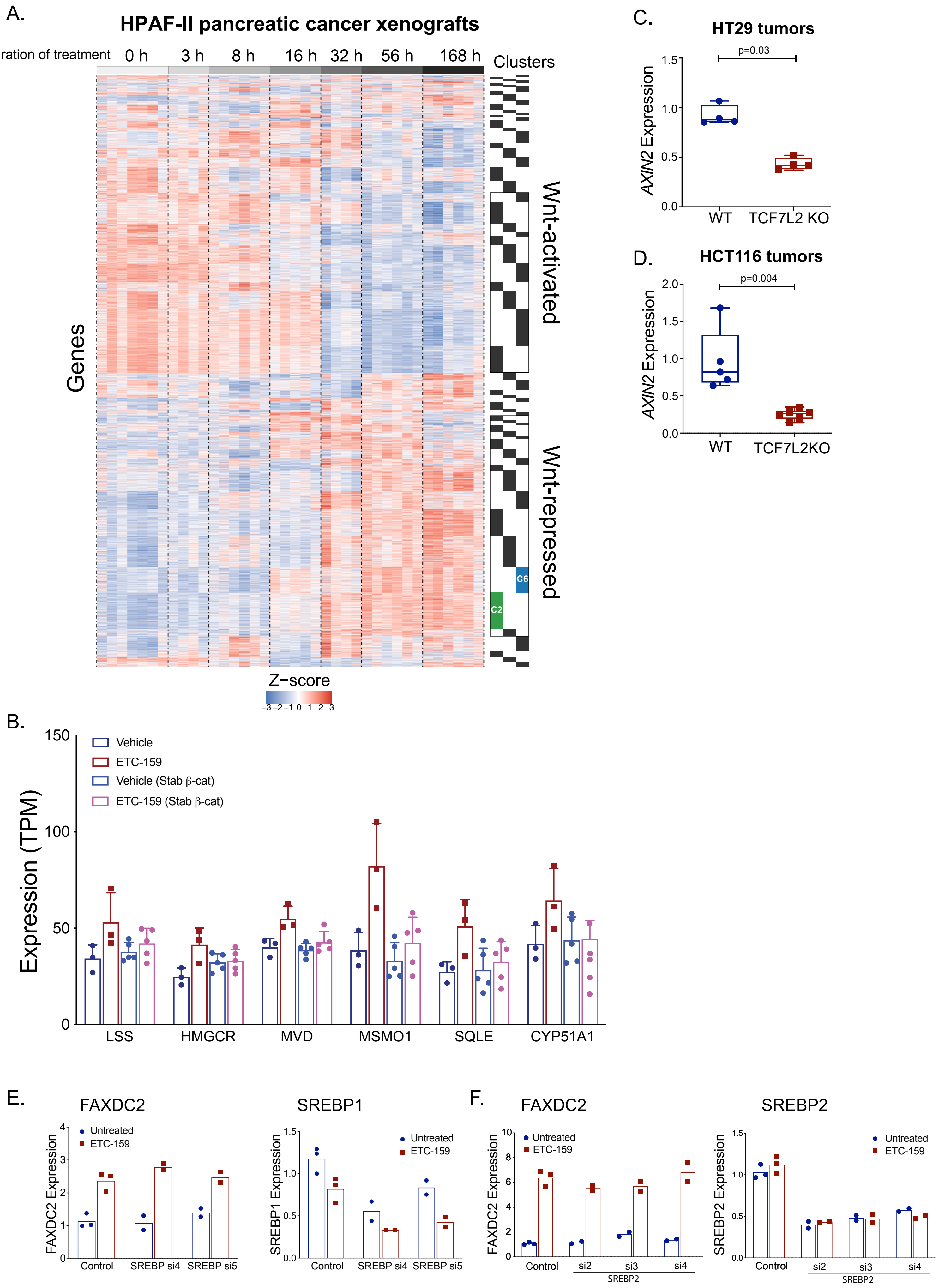
A.
HPAF-II pancreatic cancer xenografts
Duration of treatment
0 h
3 h
8 h
16 h
32 h
56 h
168 h
Clusters
Genes
Wnt-activated
Wnt-repressed
C6
C2
Z−score
−3 −2 −1 0 1 2 3
B.
Expression (TPM)
Vehicle
ETC-159
Vehicle (Stab β-cat)
ETC-159 (Stab β-cat)
LSS
HMGCR
MVD
MSMO1
SQLE
CYP51A1
C.
HT29 tumors
p=0.03
AXIN2 Expression
WT
TCF7L2 KO
D.
HCT116 tumors
p=0.004
AXIN2 Expression
WT
TCF7L2KO
E.
FAXDC2
FAXDC2 Expression
Untreated
ETC-159
Control
SREBP si4
SREBP si5
SREBP1
SREBP1 Expression
F.
FAXDC2
si2
si3
si4
SREBP2
SREBP2
SREBP2 Expression

**Figure S1:**

**A.** ***Replot of heat map of Wnt regulated genes in HPAF-II orthotopic xenografts***: Mice bearing HPAF-II orthotopic tumors were treated with a Wnt inhibitor (ETC-159, 37.5 mg twice daily). Tumors were resected at indicated time points after the treatment. RNA sequencing was performed to measure the transcript abundance. Based on the temporal dynamics of the transcriptional response to Wnt inhibition, these genes were distributed amongst 17 clusters.

**B.** ***Expression of multiple cholesterol biosynthesis pathway genes is Wnt/β-catenin dependent.*** Expression of indicated sterol pathway genes in HPAF-II orthotopic xenografts with or without stabilized β-catenin from mice treated with vehicle or ETC-159 for 56 hours was analyzed by RNA sequencing. Data shows the relative gene expression in the four groups. N=3-5 mice/group.

**C-D.** ***TCF7L2 knockout increases Axin2 expression:*** *TCF7L2* knockout HT-29 and HCT116 colon cancer xenografts had lower levels of *AXIN2* compared to the WT controls.

**E-F.** ***SREBP knockdown does not alter Wnt-regulated FAXDC2 expression***. HPAF-II cells were transfected with multiple siRNAs against *SREBP1 or SREBP2* for 24 h, followed by treatment with ETC-159 for 72 h. Expression of *FAXDC2* and *SREBP* was measured by qRT-PCR.

### Figure S2

A.

```
HS FAXDC2      1    MkgeAGHMLHNEKS[24]VAFWNSV   TWHLQ[5]SGYFWQAQWERLLTTFegKEWILFF-IGAIQVPCLFFWSFNGLL  98
AmpQe W3       1    MkfcVAAPLEATK-[13]VAISNSL   QWHME[5]SKAFFVEKWAYIYYNFgyENFFVMYiLGTSIYGLVVYTFINMLF  87
AT SMO21       1    M---DS--------    -------   -----   ---LVESGWKYLVTHF--SDFQLAC-IGSFILHESVFFLSGLPY  41
AT SMO22       1    M---AS--------    -------   -----   ---FVESGWQYLVTHF--SDFQLAC-IGSFLLHESVFFLSGLPF  41
AmpQe X4       1    M---PL---N----    IFGSALY[5]VGNIT[3]DDYLLEPSWKYMTTNY--SRFTISF-WFSVIIHEISYFGLCLPG  65
HS MSMO1       1    M---AT---NESVS    IFSSASL[4]VDSLL[5]QEPF-KNAWNYMLNNY--TKFQIAT-WGSLIVHEALYFLFCLPG  69
AT SMO11       1    M---IPYATVEEAS    IALGRNL   -----   --TRLETLWFDYSATK--SDYYLYC--HNILFLFLVFSLVPLPL  56
AT SMO13       1    M---IPYPTVEDAS    VALGRNL   -----   --TWFETVWFDYSATK--SNFHVYC--HTILVLFLVFSLAPFPL  56
AT SMO12       1    M---IPYATIEEAS    IALSRNL   -----   --TWLETLWFDYSATK--SDYYLYC--HNILFLFLIFSLVPLPL  56
YSCERG25       1    Ms-----AVFNNAT[ 7]STYSQTL[2]VAHYQ[1]QLNFMEKYWAAWYSYM--NNDVLATgLMFFLLHEFMYFFRCLPW  73

HS FAXDC2     99    LVVDTT-gkPNFISRYRIQVGKNEPVDpvKLRQSIRTVLFNQCMISFPMVVFLYPFLKWWRDPCRRE-LPTFHWFLLELA  176
AmpQe W3      88    GFVDVT-grPQIFKKYKIQDTKNFPVDpaKYKKCLQVVTFNSLLIGPLFLVVSSPIAYWRGLNCGYQ-LPTFPQVICQLI  165
AT SMO21      42    IFLER----TGFLSNYKIQT-KSNTPE--AQGKCIARLLLYHCCVNLPLMMASYPVFRFMGMESSFP-LPSWKVVSAQIL  113
AT SMO22      42    IFLER----QGFLSKYKIQT-KNNTPA--AQGKCITRLLLYHFSVNLPLMLASYPVFRAMGMRSSFP-LPSWKEVSAQIL  113
AmpQe X4      66    FLAQF----LPFMQKYKIQREKPESVD--GQWKCFLQLLFNHFIIQAPMISGLFYYNEIMGVQYDWEsMPRWYMLFFNCF  139
HS MSMO1      70    FLFQF----IPYMKKYKIQKDKPETWE--NQWKCFKVLLFNHFCIQLPLICGTYYFTEYFNIPYDWErMPRWYFLLARCF  143
AT SMO11      57    VFVELArsaSGLFNRYKIQPKVNYSLS--DMFKCYKDVMTMFILVVGPLQLVSYPSIQMIEIRSGLP-LPTITEMLSQLV  133
AT SMO13      57    VIVEW----TGWFDQFKIQKKVKYSLS--DMFQCYKEVMKLFLLVVGTLQIVSYPSIQMVGIRSGLP-LPSLMEIVAQLV  129
AT SMO12      57    VFIESSqstSDLFNRYKIQPKVKNSFS--SMFKCYKDVMKMFILVVGPLQLVSYPSIQMIEIRSGLP-LPSCMEIVAQLV  133
YSCERG25      74    FIIDQI----PYFRRWKLQPTKIPSAK--EQLYCLKSVLLSHFLVEAIPIWTFHPMCEKLGITVEVP-FPSLKTMALEIG  146

HS FAXDC2    177    IFTLIEEVLFYYSHRLLHHPTFYKKIHKKHHEWTAPIGVISLYAHPIEHA[2]-NMLPVIVGPLVMGSHLSSITMWFSLA  254
AmpQe W3     166    VFTVSVETGFYYMHRLFHHRSLYSRIHKIHHEWTAPISLASVYCHPIEHF[5]PIMLGPIILGTWFSNHLSAVWLWVAIA  247
AT SMO21     114    FYFIIEDFVFYWGHRILHTKWLYKNVHSVHHEYATPFGLTSEYAHPAEIL   FLGFATIVGPALTGPHLITLWLWMMLR  190
AT SMO22     114    FYFIIEDFVFYWGHRILHSKWLYKNVHSVHHEYATPFGLTSEYAHPAEIL   FLGFATIVGPALTGPHLITLWLWMVLR  190
AmpQe X4     140    LCLVIEDTWHYFIHQLLHHRSIYKYVHKVHHHYQAPFGMVAEYAHPIETL   VLGAGFFIGVLLFCNHVVFMWLWMFVR  216
HS MSMO1     144    GCAVIEDTWHYFLHRLLHHKRIYKYIHKVHHEFQAPFGMEAEYAHPLETL   ILGTGFFIGIVLLCDHVILLWAWVTIR  220
AT SMO11     134    VYFLIEDYTNYWVHRFFHSKWGYDKIHRVHHEYTAPIGYAAPYAHWAEVL   LLGIPTFMGPAIAPGHMITFWLWIALR  210
AT SMO13     130    VYFLIEDYTNYWIHRWMHCKWGYEKIHRIHHEYTSPIGYASPYAHWAEIL   ILGIPTFLGPAIAPGHIMTFWLWISLR  206
AT SMO12     134    VYFLVEDYTNYWVHRFFHCKWGYEKFHHIHHEYTAPIGYAAPYAHWAEVL   LLGIPTFLGPAIAPGHMITFWLWIALR  210
YSCERG25     147    LFFVLEDTWHYWAHRLFHYGVFYKYIHKQHHRYAAPFGLSAEYAHPAETL[5]TVGMPILYVMYTGKLHLFTLCVWITLR  228

HS FAXDC2    255    LIITTISHCGYHLPFLPS      --------PEFHDYHHLKFNQCYG----VLGVLDHLHGTDTMFKQTKAYERhVLLLGF  318
AmpQe W3     248    IVNTTFSHCGYHLPFLSS      --------PEGHDFHHSKFNQNFG----VLGILDRLHGTDNVFVNSIEHKRhFILLGL  311
AT SMO21     191    VIETVEAHCGYHFPWSPS[12]MWESFAYSADFHDYHHRLLYTKSGNYSSTFVYMDWIFGTDKGYRKLKAL-----KET-  272
AT SMO22     191    VLETVEAHCGYHFPWSLS[ 8]--------ADFHDYHHRLLYTKSGNYSSTFVYMDWIFGTDKGYRRLKTL-----KENG  261
AmpQe X4     217    LLETIEVHSGYDFPYLNP[ 1]NLIPGYAGVRFHDFHHKNF---NGNYSSSFRWWDWLFGTDRQYKEFVAA-----QEEV  285
HS MSMO1     221    LLETIDVHSGYDIP-LNP[ 1]NLIPFYAGSRHHDFHHMNF---IGNYASTFTWWDRIFGTDSQYNAYNEK-----RKKF  288
AT SMO11     211    QMEAIETHSGYDFPWSPT      KYIPFYGGAEYHDYHHYVGGQSQSNFASVFTYCDYIYGTDKGYRFQKKLLE-QIKESS  285
AT SMO13     207    QFEAIETHSGYDFPWSVT      KLIPFYGGPEYHDYHHYVGGQSQSNFASVFTYCDYIYGTDKGYRIHKKLLHhQIKEEA  282
AT SMO12     211    QIEAIETHSGYDFPWSLT      KYIPFYGGAEYHDYHHYVGGQSQSNFASVFTYCDYIYGTDKGYRFQKKLLQ-QMKEKS  285
YSCERG25     229    LFQAVDSHSGYDFPWSLN      KIMPFWAGAEHHDLHHHYF---IGNYASSFRWWDYCLDTESGPEAKASREE-RMKKRA  300

HS FAXDC2    319    TPLSES[5]KR-mE   333
AmpQe W3     312    SSAKEL[5]KMamD   327
AT SMO21            ------   -----
AT SMO22     262    DMKQT-   -----   266
AmpQe X4     286    KKKE--   -----   289
HS MSMO1     289    EKKTE-   -----   293
AT SMO11     286    KKSNKH[4]KS--D   298
AT SMO13     283    EEKRVR   KH--D   291
AT SMO12     286    KKSNKL[5]KF--D   299
YSCERG25     301    ENNAQK   KT--N   309
```

EV all primaries

B.

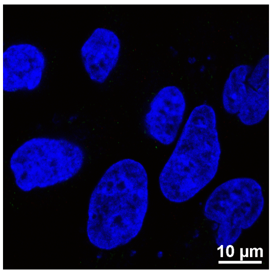

#### Figure S2 (accompanying Figure 2)

**A.** Sequence alignment comparing FAXDC2 with C4-methyl sterol oxidase enzymes from humans, plants, fungi, and predicted proteins from a sponge. The protein sequence of human HS FAXDC2 (Q96IV6) was aligned using the NCBI COBALT Multiple Alignment tool (17) against human HsMSMO1 (Q15800), *Saccharomyces cerevisiae* Erg25/YSCERG25 (P53045), *Arabidopsis thaliana* SMO11, 12, 13, 21 and 22 (Q8L7W5, Q1EC69, F4JLZ6, Q9ZW22, and Q8VWZ8 and predicted proteins from *Amphimedon queenslandica* (AmpQe) (18), XP_003386944 (W3) and XP_003385270.3 (X4). Identical residues are highlighted in red, and the three highly conserved histidine domains are boxed.

**B.** Antibody control for figure 2D. HeLa cells were co-transfected with vectors encoding epitope-tagged constructs of MSMO1, NSDHL, and FAXDC2. Cells visualized after staining with fluorescence-tagged secondary antibodies show no background staining.

Figure S3

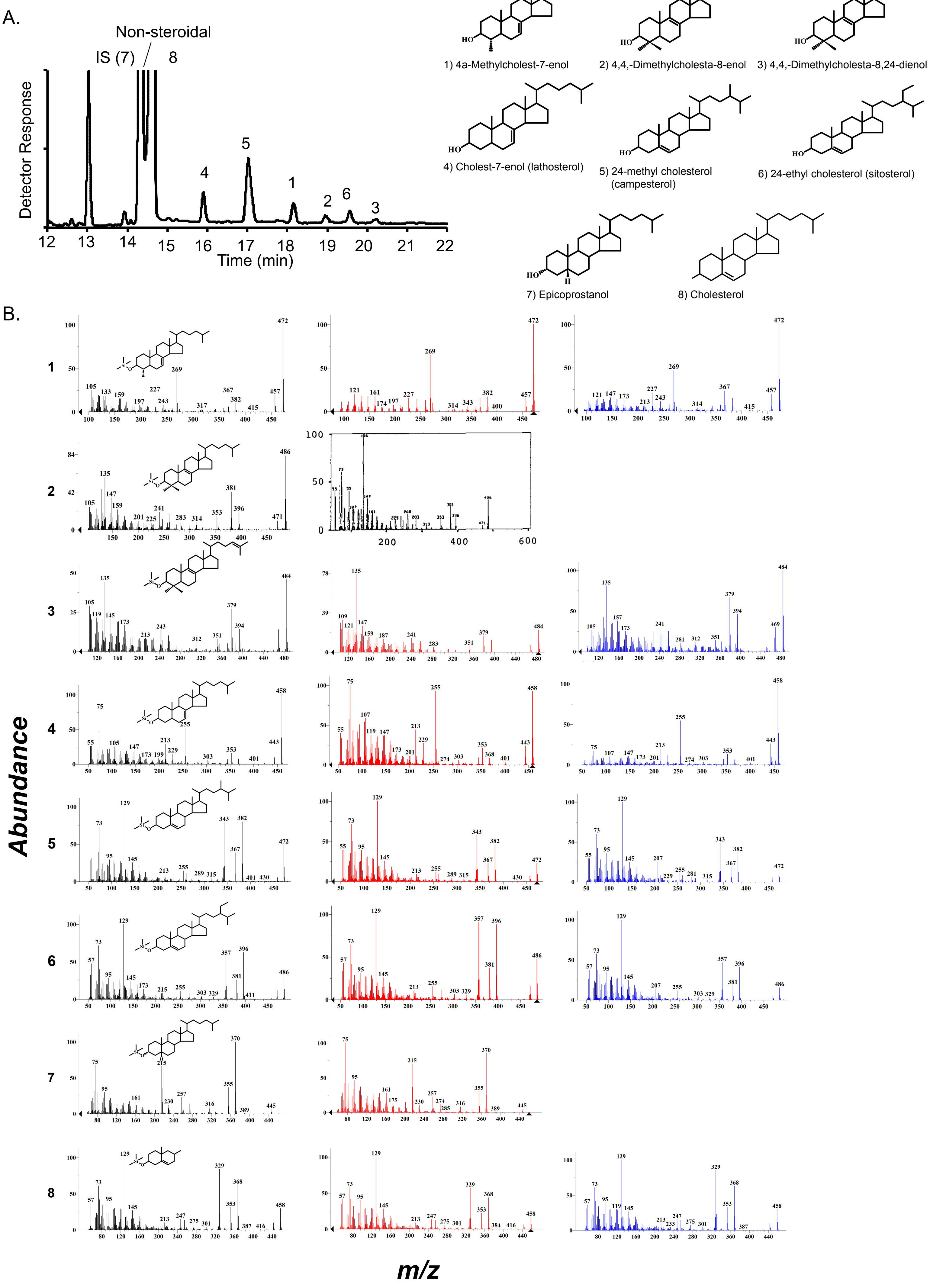

A.
Non-steroidal
IS (7)
8
Detector Response
Time (min)
1) 4a-Methylcholest-7-enol
2) 4,4,-Dimethylcholesta-8-enol
3) 4,4,-Dimethylcholesta-8,24-dienol
4) Cholest-7-enol (lathosterol)
5) 24-methyl cholesterol (campesterol)
6) 24-ethyl cholesterol (sitosterol)
7) Epicoprostanol
8) Cholesterol
B.
Abundance
m/z

**Figure S3: Identification of methyl sterols using GC-MS (accompanying Figure 2)**

**A.** Representative GC trace of total sterols recovered from a representative HPAF-II tumor. Internal standard (IS) epicoprostanol (3α-inverted C-OH and A/B *cis* ring generating a bent ring structure affording a sterol standard that elutes before cholesterol and does not interfere with the sterol analysis of natural metabolites) and sterols identified in the chromatogram are indicated.

**B.** Mass spectra of TMS-derivatized sterols isolated from tumors and corresponding spectra of standards. Mass spectra in the left column are from tumor samples, the middle column is from the NIST library (red) or from (6) (black), and the right column (blue) from the Nes sterol collection.

Figure S4

A.

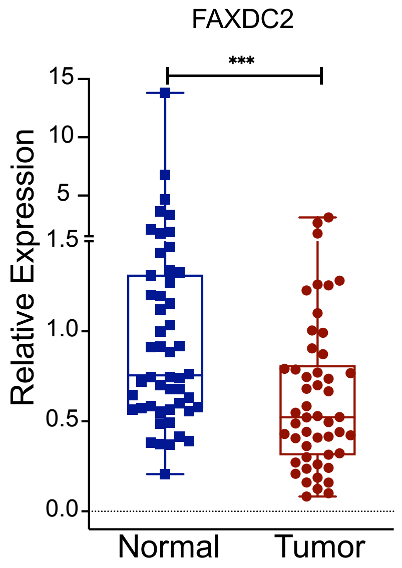


B. Sequence of the FAXDC KO clone # 3

```
TGCACCTGGCCTCAGTTTCCTTGTTGATCAGTGAGCCCTAGACAGGATGAGGAGCCA-------------------------------------------

tgcacctggcctcagtttccttgttgatcagtgagccctagacaggatgaggagccagctgtgcctcaggcagggcctgctcacCGGTGTGAATAGTAGA
                                                                                   <----------------
                                                                                        Exon 5
-----------------------------------TCCA---------------------------CAC-------------------------------
                                   ||||                           |||
ACAAGACTTCCTCGATCAGCGTGAAGATGGCCAGCTCCAGGAGGAACCAGTGGAAGGTGGGTAGCTCACGGCGGCAGGGGTCTCTCCACCATTTGAGGAA
<---------------------------------------------------------------------------------------------------
        Exon 5
-----------------------------------------------AAAGAACTGTGTGGATAGACTGGCGCAGTTTCACAGGATCCACCTGCCCAAGG
                                                           |||||||||||||||||||||||||||||||||||||||||
GGGATAGAGGAAGACCACCATGGGGAAAGATATCATGCACTGGTTGAAAAGAACTGTGCGGATAGACTGGCGCAGTTTCACAGGATCCACctgcccaagg
<--------------------------------------------------------------------------------------------
                      Exon 5
```

C. Sequence of the FAXDC KO clone #12

```
gGTGGATCCTGTGAAACTGCGCCAGTCTATCCGCACAGTTCTTTTCAACCAGTGCATGATATCTTTCCCCATGGTGGTCTTCCTCTATCCCTTCCTCAAATGGTG
|||||||||||||||||||||||||||||||||||||||||||||||||||||||||||||||||||||||||||||||||||||| |
gGTGGATCCTGTGAAACTGCGCCAGTCTATCCGCACAGTTCTTTTCAACCAGTGCATGATATCTTTCCCCATGGTGGTCTTCCTCT-T-----------------
 -------------------------------------------------------------------------------------------------------->
                                                 Exon 5
CCCTTCCTCAAATGGTGGAGAGACCCCTGCCGCCGTGAGCTACCCACCTTCCACTGGTTCCTCCTGGAGCTGGCCATCTTCACGCTGATCGAGGAAGTCTTGTTC
                 ||| ||              ||  | |||  |
-----------------GAG-GA--------------AG--A-CCA--T--------------------------------------------------------
--------------------------------------------------------------------------------------------------------->
                                                 Exon 5
TACTATTCACACCGgtgagcaggccctgcctgaggcacagctggctcctcatcctgtctagggctcactgatcaacaaggaaactgaggccaggtgcagtgact

-------------------------------------------------------------------------------------------------------
------------->
cacgcttgtaatcctggcaatttgggaggctgaggtgggcggatcacttgagcccaggagtttgagac
                                 |||||||||||||||||||||||||||||||||||
---------------------------------ggtgggcggatcacttgagcccaggagtttgagac
```

D.

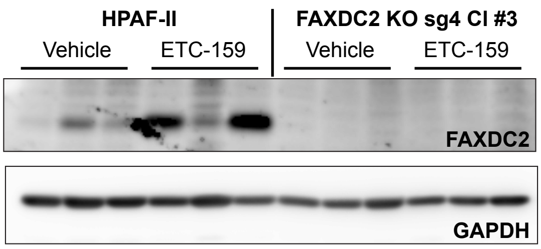

#### Figure S4

**A.** **FAXDC2 is repressed in primary colorectal cancers compared to adjacent normal tissues:** Expression of FAXDC2 was compared in primary cancers and adjacent normal tissues by qRT-PCR.

**B-C.** ***FAXDC KO clones obtained using CRISPR methodology show the deletion of large regions of Exon 5***. The genomic sequence of *FAXDC2* knockout clone 3 (A) and clone 12 (B) both show deletion of a large region of Exon 5.

**D.** ***Loss of FAXDC2 protein in the FAXDC2 KO xenografts*.** Protein lysates from the parental HPAF-II and FAXDC2 KO xenografts from the vehicle or ETC-159 treated mice were probed with the FAXDC2 or GAPDH antibody. Each lane represents tumor lysate from an individual mouse.

Figure S5

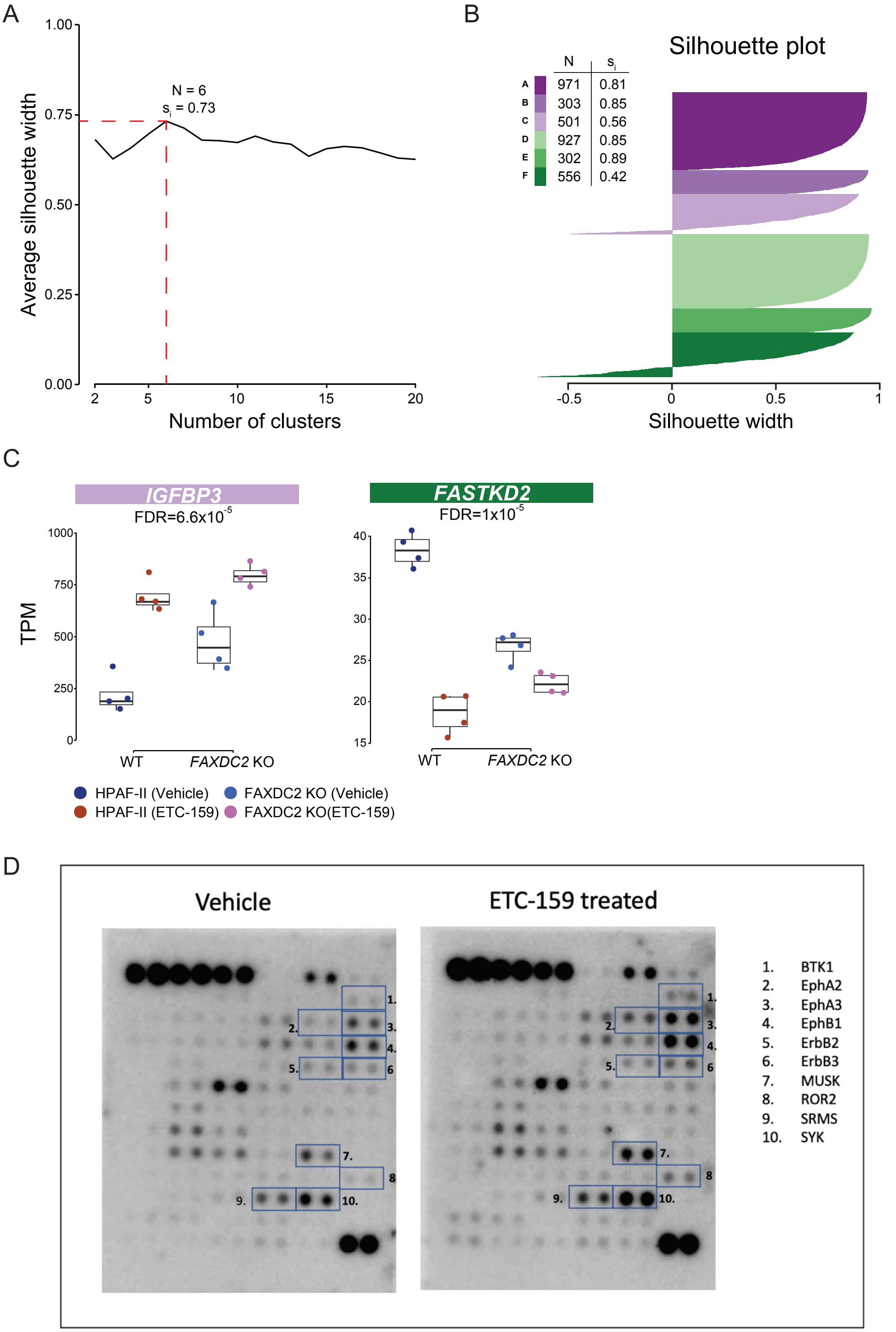
A
1.00
0.75
0.50
0.25
0.00
Average silhouette width
N = 6
$s_i$ = 0.73
2
5
10
15
20
Number of clusters
B
Silhouette plot
N
$s_i$
A 971 0.81
B 303 0.85
C 501 0.56
D 927 0.85
E 302 0.89
F 556 0.42
-0.5
0
0.5
1
Silhouette width
C
IGFBP3
FDR=6.6x10$^{-5}$
FASTKD2
FDR=1x10$^{-5}$
TPM
1000
750
500
250
0
40
35
30
25
20
15
WT
FAXDC2 KO
HPAF-II (Vehicle)
FAXDC2 KO (Vehicle)
HPAF-II (ETC-159)
FAXDC2 KO(ETC-159)
D
Vehicle
ETC-159 treated
1. BTK1
2. EphA2
3. EphA3
4. EphB1
5. ErbB2
6. ErbB3
7. MUSK
8. ROR2
9. SRMS
10. SYK

#### Figure S5

**A.** FAXDC2-dependent genes form six distinct clusters. Clustering FAXDC2-dependent genes (FDR < 10%) and calculating the average silhouette width for different clusters find six clusters as the optimal number.

**B.** A silhouette plot for the FAXDC2-dependent genes clustered into six distinct clusters reveals that most genes are well clustered.

**C.** Representative genes from 2 clusters of FAXDC2-dependent genes (TPM - transcripts per million).

**D.** ***Increased phosphorylation of multiple kinases in HPAF-II xenografts treated with a Wnt-inhibitor*:** Tumor lysates from orthotopic xenografts from mice treated with vehicle or ETC-159 were analyzed using phosphotyrosine arrays. Proteins with >1.5 fold increase in density are indicated.

Figure S6

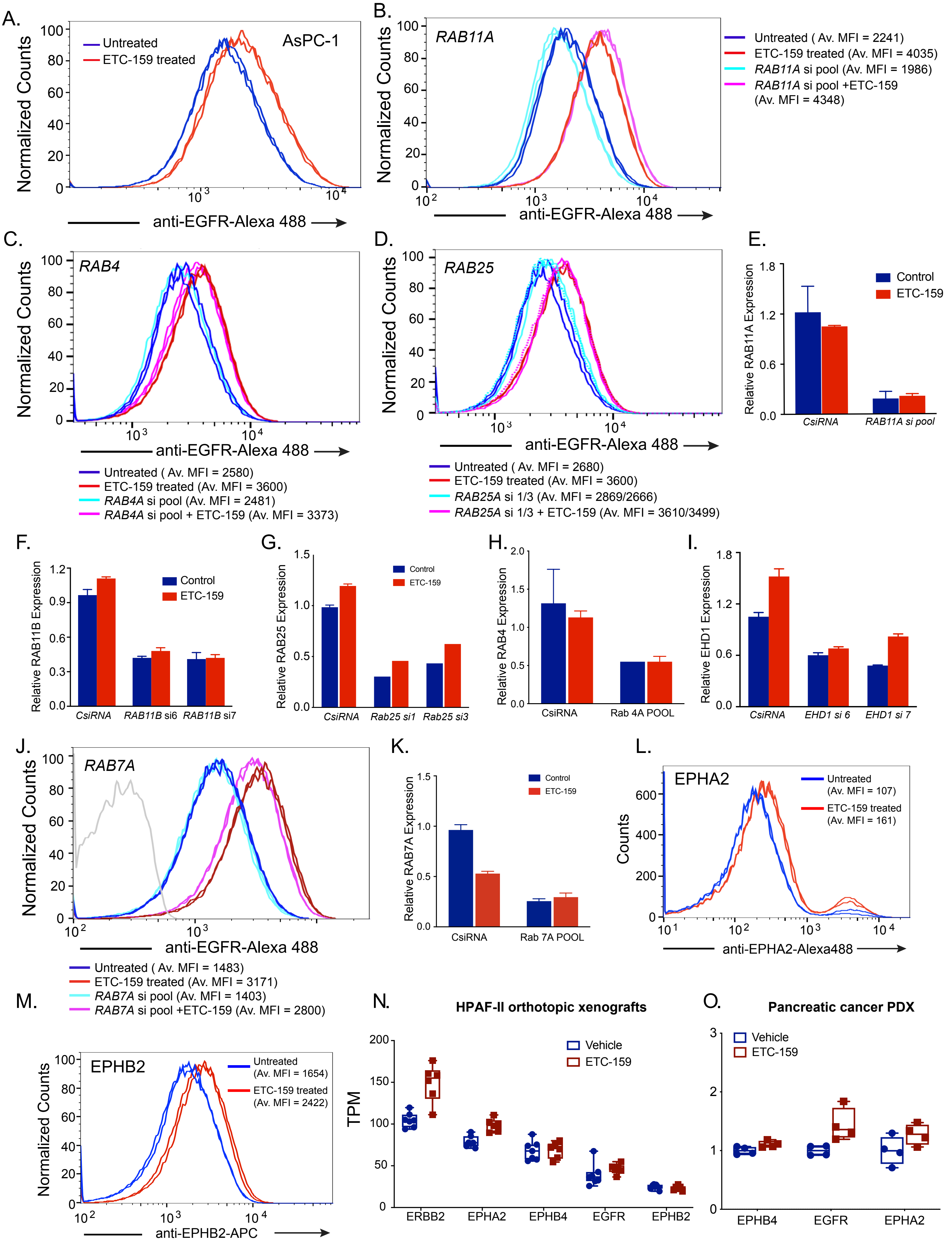
A.
AsPC-1
Untreated
ETC-159 treated
Normalized Counts
anti-EGFR-Alexa 488
B.
RAB11A
Untreated ( Av. MFI = 2241)
ETC-159 treated (Av. MFI = 4035)
RAB11A si pool (Av. MFI = 1986)
RAB11A si pool +ETC-159 (Av. MFI = 4348)
C.
RAB4
Untreated ( Av. MFI = 2580)
ETC-159 treated (Av. MFI = 3600)
RAB4A si pool (Av. MFI = 2481)
RAB4A si pool + ETC-159 (Av. MFI = 3373)
D.
RAB25
Untreated ( Av. MFI = 2680)
ETC-159 treated (Av. MFI = 3600)
RAB25A si 1/3 (Av. MFI = 2869/2666)
RAB25A si 1/3 + ETC-159 (Av. MFI = 3610/3499)
E.
Relative RAB11A Expression
Control
ETC-159
CsiRNA
RAB11A si pool
F.
Relative RAB11B Expression
RAB11B si6
RAB11B si7
G.
Relative RAB25 Expression
Rab25 si1
Rab25 si3
H.
Relative RAB4 Expression
Rab 4A POOL
I.
Relative EHD1 Expression
EHD1 si 6
EHD1 si 7
J.
RAB7A
Untreated ( Av. MFI = 1483)
ETC-159 treated (Av. MFI = 3171)
RAB7A si pool (Av. MFI = 1403)
RAB7A si pool +ETC-159 (Av. MFI = 2800)
K.
Relative RAB7A Expression
Rab 7A POOL
L.
EPHA2
Counts
Untreated (Av. MFI = 107)
ETC-159 treated (Av. MFI = 161)
anti-EPHA2-Alexa488
M.
EPHB2
Untreated (Av. MFI = 1654)
ETC-159 treated (Av. MFI = 2422)
anti-EPHB2-APC
N.
HPAF-II orthotopic xenografts
TPM
Vehicle
ETC-159
ERBB2
EPHA2
EPHB4
EGFR
EPHB2
O.
Pancreatic cancer PDX
EPHB4
EGFR
EPHA2

#### Figure S6 (accompanying Figure 5)

**A.** ***Wnt inhibition with ETC-159 treatment increased EGFR abundance on the cell surface***. AsPC-1 cells were treated with 100 nM ETC-159 for 72 h before analysis of EGFR levels by flow cytometry. Av. MFI = Average median fluorescence intensity of the technical replicates from the same experiment. Each histogram represents ~50,000 cells. Data is representative of three independent experiments ($p = 0.009$).

**B-D.** ***RAB11A, RAB4, or RAB25 knockdown do not alter Wnt-regulated increase in EGFR levels on the surface of HPAF-II cells***. HPAF-II cells were transfected with either a pool of siRNAs or individual siRNAs against indicated *RABs* for 24 h, followed by treatment with ETC-159 for 72 h. EGFR levels were assessed by flow cytometry as described in A.

**E-I.** Transcript levels of *RAB11,* ***RAB4, RAB25*** or EHD1 are reduced after transfection of HPAF-II cells with indicated siRNAs. Error bars represent mean ± SD.

**J.** ***RAB7A knockdown does not alter the Wnt-regulated increase of EGFR levels on the cell surface***. HPAF-II cells were transfected with a pool of siRNAs against *RAB7* for 24 h, followed by treatment with ETC-159 for 72 h. EGFR levels were assessed by flow cytometry as described in A.

**K.** Transfection of HPAF-II cells with *RAB7A siRNA* reduced its transcript levels. Error bars represent mean ± SD.

**L-M.** ***Wnt inhibition increases the abundance of EPHA2 and EPHB2.*** HPAF-II cells were treated with 100 nM ETC-159 for 72h before flow cytometric analysis. Av. MFI = Average median fluorescence intensity of the technical replicates from the same experiment.

**N-O.** ***Wnt inhibition does not increase transcript levels of RTKs***. Transcripts of indicated RTKs in HPAF-II xenografts or pancreatic patient-derived xenograft PAXF1861 from vehicle or ETC-159 treated mice were measured using RNAseq. Each data point represents an individual mouse, n = 4-6/group.

Figure S7

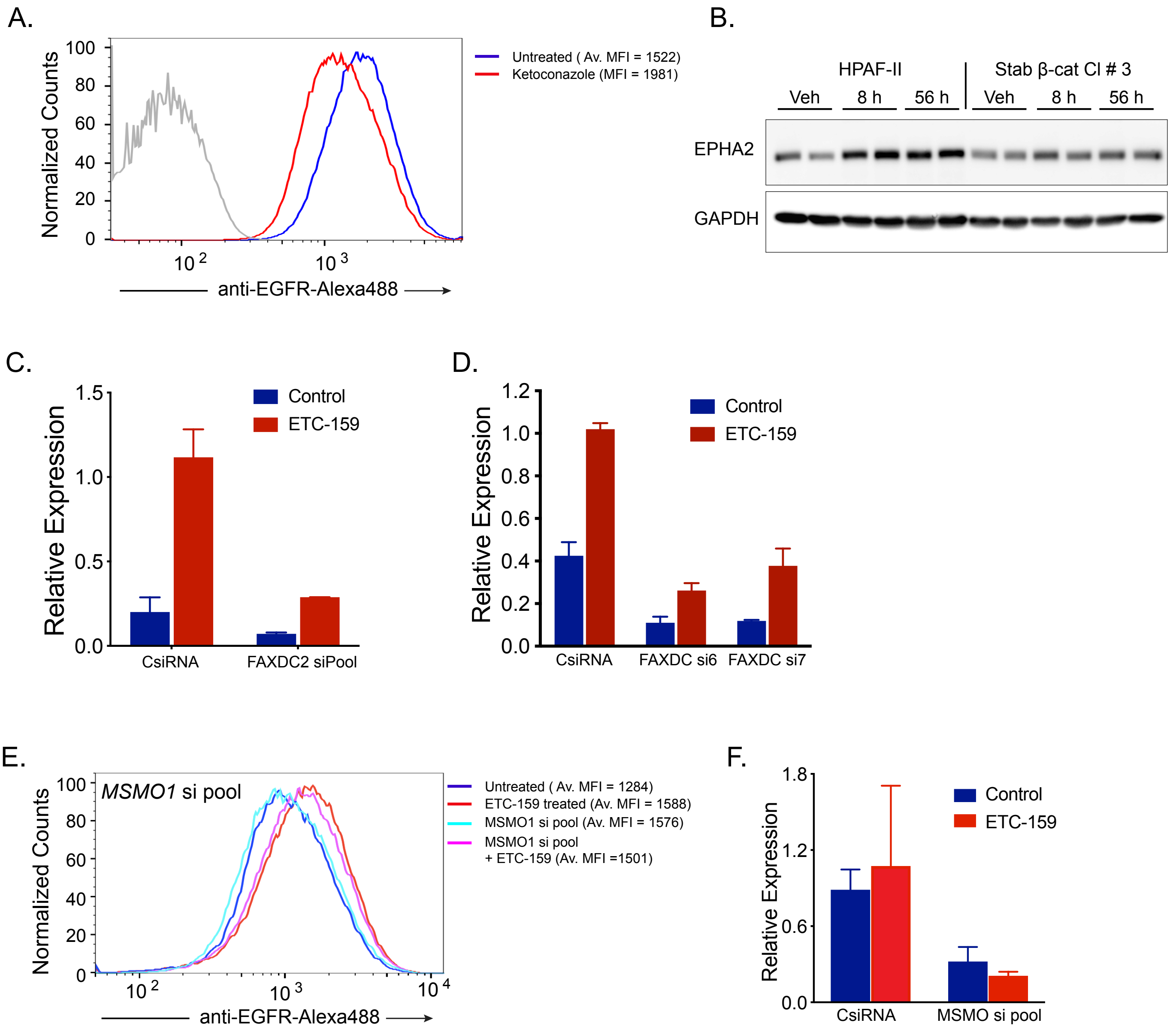
A.
Normalized Counts
100
80
60
40
20
0
10 2
10 3
anti-EGFR-Alexa488
Untreated ( Av. MFI = 1522)
Ketoconazole (MFI = 1981)
B.
HPAF-II
Stab β-cat Cl # 3
Veh
8 h
56 h
Veh
8 h
56 h
EPHA2
GAPDH
C.
Relative Expression
1.5
1.0
0.5
0.0
Control
ETC-159
CsiRNA
FAXDC2 siPool
D.
Relative Expression
1.2
1.0
0.8
0.6
0.4
0.2
0.0
Control
ETC-159
CsiRNA
FAXDC si6
FAXDC si7
E.
Normalized Counts
MSMO1 si pool
100
80
60
40
20
0
10 2
10 3
10 4
anti-EGFR-Alexa488
Untreated ( Av. MFI = 1284)
ETC-159 treated (Av. MFI = 1588)
MSMO1 si pool (Av. MFI = 1576)
MSMO1 si pool
+ ETC-159 (Av. MFI =1501)
F.
Relative Expression
1.8
1.2
0.6
0.0
Control
ETC-159
CsiRNA
MSMO si pool

**Figure S7 (accompanying Figure 6)**

**A.** ***Ketoconazole treatment increases EGFR levels on HPAF-II cells***. HPAF-II cells were treated with 6 µg/mL ketoconazole for 72 h before analysis of EGFR by flow cytometry. Data is representative of three independent experiments. Each histogram represents ~50,000 cells.

**B.** ***Stabilized β-catenin prevents Wnt inhibition-mediated increase in EPHA2 levels***. Protein lysates from HPAF-II xenografts with stabilized β-catenin from mice treated with vehicle or ETC-159 were immunoblotted for EPHA2. Each lane represents tumor lysate from an individual mouse.

**C-D.** ***ETC-159 treatment of the HPAF-II cells increases FAXDC2 transcript levels.*** Transfection of HPAF-II cells with *FAXDC2* siRNA pool or individual siRNAs reduce its transcript levels. Error bars represent mean ± SD.

**E.** ***MSMO1 knockdown does not affect EGFR abundance on ETC-159 treated cells.*** HPAF-II cells were transfected with a pool of four siRNAs against *MSMO1* for 24 h, followed by treatment with ETC-159 for 72 h. Cells were stained with Alexa fluor-488 conjugated anti-EGFR antibody and analyzed by flow cytometry. Data is representative of three independent experiments, and one replicate is shown. Av. MFI = Average median fluorescence intensity of the technical replicates from the same experiment.

**F.** ***MSMO1 transcript levels in the HPAF-II cells are reduced following transfection with a pool of MSMO1 siRNAs.*** Error bars represent mean ± SD.

Figure S8

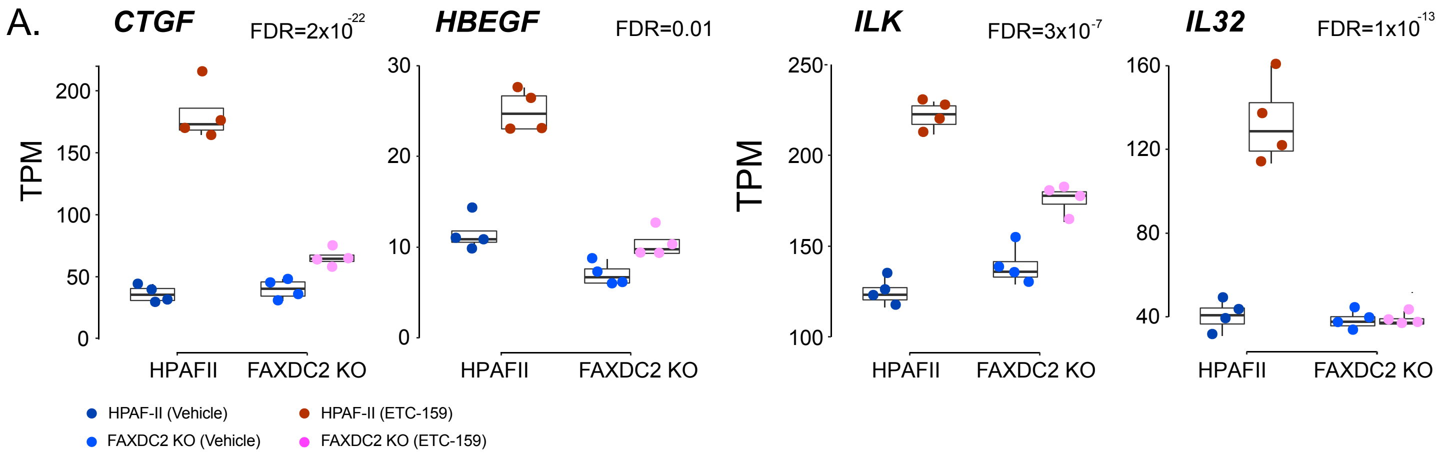

A.
CTGF
FDR=2x10^-22
HBEGF
FDR=0.01
ILK
FDR=3x10^-7
IL32
FDR=1x10^-13
TPM
HPAFII
FAXDC2 KO
HPAF-II (Vehicle)
HPAF-II (ETC-159)
FAXDC2 KO (Vehicle)
FAXDC2 KO (ETC-159)


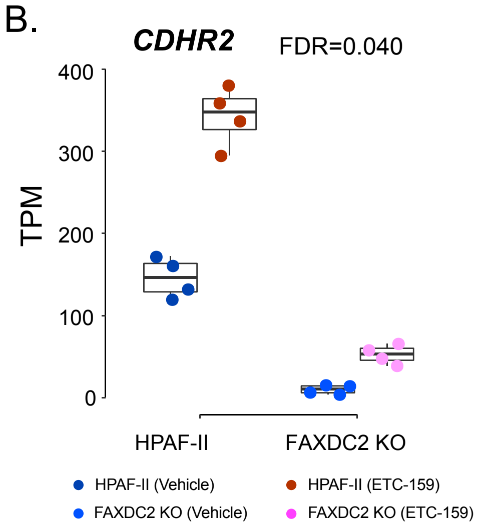

B.
CDHR2
FDR=0.040
TPM
HPAF-II
FAXDC2 KO
HPAF-II (Vehicle)
HPAF-II (ETC-159)
FAXDC2 KO (Vehicle)
FAXDC2 KO (ETC-159)


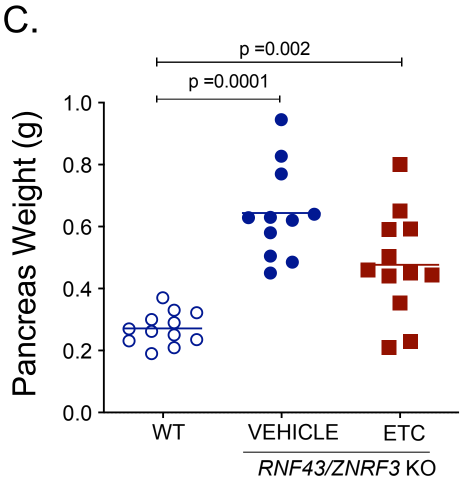

C.
p =0.002
p =0.0001
Pancreas Weight (g)
WT
VEHICLE
ETC
RNF43/ZNRF3 KO

#### Figure S8 (accompanying Figure 7)

**A.** ***FAXDC2 knockout blunts the Wnt-regulated senescence response***. The expression of selected senescence-associated genes was analyzed in the tumors of all four groups. Each data point represents an individual tumor. ETC-159 treatment increased the expression of senescence-associated genes in the HPAF-II tumors, however, no such increase was observed in the FAXDC2 knockout tumors.

**B.** ***FAXDC2 knockout blunts the Wnt-inhibition-mediated increase in cellular differentiation***. The expression of differentiation-associated genes was analyzed in the tumors of all four groups. Each data point represents an individual tumor.

**C.** ***$Ptf1a^{Cre}Rnf43^{fl/fl}/Znrf3^{fl/fl}$ mice with activated Wnt signaling have increased pancreatic weight compared to the WT mice.*** Treatment with ETC-159 for 21 days reduces this increase in pancreatic weight. Each dot represents an individual mouse. Data is from two independent biological experiments with 5-6 mice/group.
